# Integrative Omics Uncovers Low Tumorous Magnesium Content as A Driver Factor of Colorectal Cancer

**DOI:** 10.1101/2024.01.22.576593

**Authors:** Rou Zhang, Meng Hu, Yu Liu, Wanmeng Li, Zhiqiang Xu, Siyu He, Ying Lu, Yanqiu Gong, Xiuxuan Wang, Shan Hai, Shuangqing Li, Shiqian Qi, Yuan Li, Yang Shu, Dan Du, Huiyuan Zhang, Heng Xu, Zongguang Zhou, Peng Lei, Hai-Ning Chen, Lunzhi Dai

## Abstract

Magnesium (Mg) deficiency is associated with increased risk and malignancy of colorectal cancer (CRC), yet the underlying mechanisms remain elusive. Here we used genomic, proteomic, and phosphoproteomic data to elucidate the impact of Mg deficiency on CRC. Genomic analysis identified 160 genes with higher mutation frequencies in Low-Mg tumors, including key driver genes such as *KMT2C* and *ERBB3*. Unexpectedly, initiation driver genes of CRC, such as *TP53* and *APC*, displayed higher mutation frequencies in High-Mg tumors. Additionally, proteomics and phosphoproteomics indicated that low tumorous Mg content may activate epithelial-mesenchymal transition (EMT) by modulating inflammation or remodeling the phosphoproteome of cancer cells. Notably, we observed a negative correlation between the phosphorylation of DBN1 at S142 (DBN1^S142p^) and Mg content. A mutation in S142 to D (DBN1^S142D^) mimicking DBN1^S142p^ upregulated MMP2 and enhanced cell migration, while treatment with MgCl_2_ reduced DBN1^S142p^, thereby reversing this phenotype. Mechanistically, Mg^2+^ attenuated the DBN1-ACTN4 interaction by decreasing DBN1^S142p^, which, in turn, enhanced the binding of ACTN4 to F-actin and promoted F-actin polymerization, ultimately reducing MMP2 expression. These findings shed new light on the crucial role of Mg deficiency in CRC progression and suggest that Mg supplementation may offer a promising preventive and therapeutic strategy for CRC.

## Introduction

Metal ions are crucial in both physiological and pathological processes within living organisms[1, 2]. Magnesium (Mg), a predominant intracellular divalent cation, is crucial for maintaining cellular homeostasis and participates in nearly all cellular processes[3, 4]. The intracellular Mg content is up to 10−30 mM, however, the concentration of free Mg^2+^ in cells is only 0.5−1.2 mM[5]. Mg^2+^ can bind to ATP, ribosomes or nucleotides as a cofactor and serves as an activator of numerous enzymes involved in glycolysis, phosphorylation events, DNA repair, DNA stabilization, and protein synthesis[6−8]. Many proteins, such as MRS2, TRPM6/7, MAGT1, SCL41A1, and CNNMs, are well-established Mg^2+^ transporters[9]. Dysregulation of these transporters may lead to Mg^2+^ imbalance and diseases.

Mg is crucial in controlling cancer initiation and progression[5, 10]. Disruptions in Mg homeostasis contribute to cancer progression by promoting proliferation, angiogenesis, and invasion of cancer cells into surrounding tissues[11−13]. Additionally, changes in Mg levels can impair immune function, compromising the body’s ability to detect and eliminate cancerous cells[14, 15]. Furthermore, Mg’s dysregulation affects chemoresistance, rendering cancer cells less susceptible to chemotherapy[16, 17]. Colorectal cancer (CRC) ranks as the third most common cause of cancer-related deaths globally. A meta-analysis of 29 studies demonstrated that increased Mg intake is linked to a reduced risk of CRC[18−21]. Previous investigations showed that Mg’s anti-inflammatory properties may help reduce inflammation in the colon[22, 23], which is a risk factor for CRC[24]. Moreover, Mg functions as an antioxidant, protecting colon cells against oxidative stress and preventing DNA damage[25−27]. In CRC treatment, Mg can enhance the effectiveness of chemotherapy by improving drug uptake by cancer cells and protecting healthy cells from damage[28].

Despite some progress, many questions about the role of Mg in CRC remain unanswered. First, there is a lack of systematic investigation on how Mg affects tumor progression at the molecular level. Second, it is unclear whether Mg has a differential effect on left– and right-sided CRC at the molecular level[29]. Third, tumor metastasis is a leading cause of mortality in CRC patients. Mg deficiency not only reduces intracellular ATP[30] but also affects kinase and phosphatase activities[31],[32]. Dysregulated protein phosphorylation is common in CRC and linked to unfavorable outcomes[33–36]. However, the direct link between Mg deficiency and tumor metastasis through affecting protein phosphorylation still lacks direct evidence.

With the rapid development of omics technologies, the functional interpretation of metal ions in aging using omics data has been achieved[37]. In this study, we utilized an integrative genomic, proteomic, and phosphoproteomic approach to uncover the roles of Mg in CRC and found that low Mg content in tumors affected both genomic stability and metastasis. Genomics revealed that low Mg content in tumors increased the gene mutations associated with tumor progression rather than initiation. Proteomics and phosphoproteomics analyses indicated that low Mg content in tumors could activate epithelial-mesenchymal transition (EMT) by activating the complement pathway and inducing inflammation or by directly remodeling protein phosphorylation in cancer cells. Furthermore, we demonstrated that Mg^2+^ weakened the interaction between DBN1 and ACTN4 by reducing the phosphorylation of DBN1 at S142 (DBN1^S142p^), which enhanced the interaction of ACTN4 to F-actin and promoted F-actin polymerization, ultimately leading to downregulated MMP2 and reduced cancer cell migration.

## Results

### Low Mg content in tumors predicts unfavorable prognosis of CRC patients

To uncover the clinical significance of low Mg content in tumors and its impact on tumor progression, omics data of 230 paired tissue samples collected from 115 treatment-naive CRC patients were used (**Figure 1**A; Table S1). Proteomics and phosphoproteomics employed a TMT-based quantitative approach (Figure S1A). Correlation analysis of quality control (QC) samples, internal standard (IS) samples in proteomics and phosphoproteomics, and replicate samples in proteomics demonstrated the stability of the instrument as well as the reliability and reproducibility of mass spectrometry (MS) data (Figure S1B−H). A total of 9652 proteins and 12,988 phosphosites were identified. Of them, 5322 proteins (with > 1 unique peptide) and 2162 phosphosites detected in > 50% of samples were utilized for further data analysis (Figure S1E, F, and S1I−L). Additionally, an inductively coupled plasma-mass spectrometry (ICP-MS) analysis of Mg content was conducted on 115 paired samples, and correlation analysis of QC samples indicated the reliability of the measurements (Figure S1M). Whole-exome sequencing (WES) data were obtained from a previous work[38], in which 16,234 mutated genes with less than 5000 amino acids in 76 paired samples were reported.

**Figure 1.**
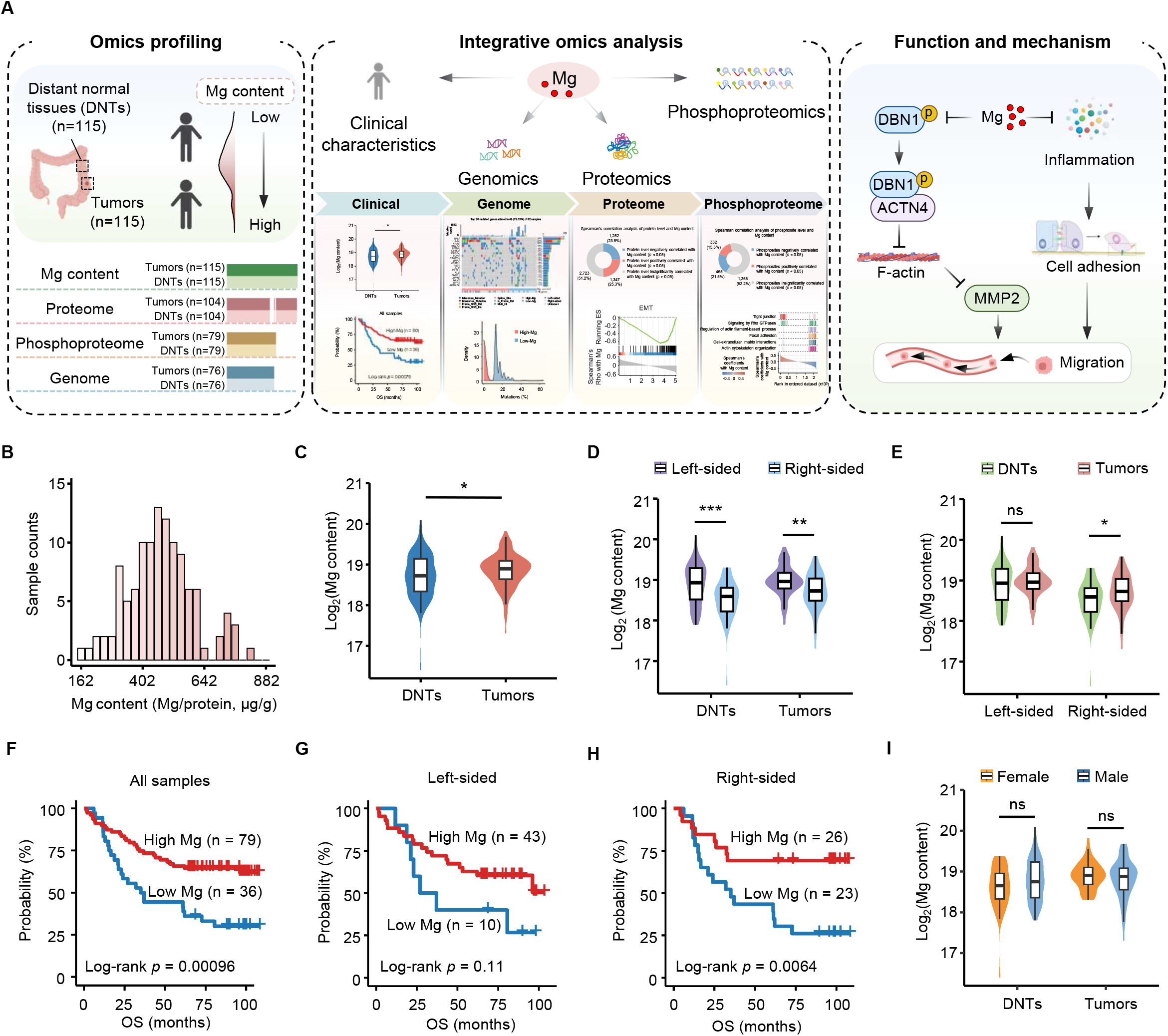
Low Mg content in tumors predicts unfavorable prognosis of CRC. **A.** The Study’s concept and workflow. **B.** Intra-tumorous Mg content of CRCs. **C.** Statistics of Mg content in the tumors and paired DNTs of 115 CRCs. *, *P* < 0.05. Wilcoxon rank-sum test. **D.** Comparisons of Mg content between left– and right-sided tumors or between left– and right-sided DNTs. ***, *P* < 0.001; **, *P* < 0.01. Wilcoxon rank-sum test. **E.** Comparisons of Mg content between tumors and paired DNTs located on the left or right side. *, *P* < 0.05; ns, means no significance. Wilcoxon rank-sum test. **F.** Survival analysis of 115 CRC patients with different Mg contents in tumors. Log-rank test. **G.** Survival analysis of 53 left-sided CRC patients with different Mg contents in tumors. Log-rank test. **H.** Survival analysis of 49 right-sided CRC patients with different Mg contents in tumors. Log-rank test. **I.** Comparisons of Mg content between females and males in tumors or DNTs. ns, means no significance. Wilcoxon rank-sum test.

Our findings showed that the intra-tumoral Mg content ranged from 162−920 μg per gram of extracted protein (Figure 1B). The Wilcoxon rank-sum test identified higher levels of Mg content in tumors than in distant normal tissues (DNTs) (Figure 1C), and the levels of Mg between the left– and right-sided tumors were significantly different (Figure 1D and E). Our Kaplan−Meier survival analysis of 115 CRC patients revealed that lower levels of Mg in tumors were associated with poor overall survival (OS) (Figure 1F). Multivariable Cox regression models demonstrated that Mg content remains significantly associated with CRC survival even after adjusting for prognostic factors, such as renal function and liver function (Figure S2A−J), indicating that Mg content serves as an independent predictor of survival. Moreover, we conducted an independent analysis to investigate the relationship between Mg levels and prognosis in left-sided and right-sided CRC cases. Interestingly, we observed a notable connection between Mg levels and the prognosis of CRC patients specifically in the right-sided cases, while a less significant association was found in the left-sided cases (Figure 1G and H). Additionally, we also did not observe any significant variation in Mg content between tumors in female and male patients (Figure 1I). Therefore, the following correlation analysis between Mg and omics data did not take into account the influence of gender.

### Low Mg content in tumors is linked to genome instability

Mg is important for DNA stabilization, DNA replication and DNA repair[39, 40]. To investigate the impact of low Mg content on CRC at the genomic level, we analyzed Mg-associated genomic data. Of the 76 colon cancer cases analyzed by WES, 20 showed hypermutation in tumors (> 10 Mut/Mb) (Table S2)[38]. To exclude hypermutations caused by mutated mismatch repair (MMR) genes and major replicase genes, including *MSH2*, *MSH6*, *MLH1*, *PSM2*, *POLD,* and *POLE*, we removed 14 hypermutated samples with mutations in the above genes[38, 41]. Statistics showed that *TP53* had the highest frequency of mutations (48%), followed by *APC* (40%) and *KRAS* (39%), in the remaining 62 colon cancer cases (**Figure 2**A). The total percentage of single nucleotide variants (SNVs) differed between the High-Mg (39 cases) and Low-Mg (23 cases) groups, with the most frequent thymine to cytosine (T > C) transition in the Low-Mg group (Figure 2B). Additionally, the classes of mutations were different between the two groups, and more types of mutations were observed in the High-Mg group (Figure 2C). Association analysis of mutated genes in the two groups using the somatic interactions algorithm showed that mutations in the High-Mg group were mutually exclusive, while mutations in the Low-Mg group mainly co-occurred (Figure 2D and E), suggesting that low Mg content may cause simultaneous mutations in many genes, possibly due to increased genome instability and the dysregulation of DNA replication and repair processes[39, 40, 42−44]. In addition, somatic copy number alteration (SCNA) analysis revealed distinct patterns in the cytobands at the locations of the variant sites. In the High-Mg group, amplified regions were observed at 5p15.33 and 8p23.1, while in the Low-Mg group, amplified regions were found at 4p16.1, 15q11.1, and 15q11.2. Conversely, deleted regions were identified at 1p36.21, 1p36.33, and 9p11.2 in the High-Mg group and at 17q12 and 17q21.31 in the Low-Mg group (Figure 2F). Among the identified 7,173 mutated genes with less than 5000 amino acids in 62 paired samples, the mutation frequencies of 162 genes were associated with Mg content, of which 160 genes exhibited a higher frequency of mutations in the Low-Mg group (Figure 2G and H; Table S2), including 6 known CRC driver genes such as *KMT2C*, *BCL9*, *ERBB3*, *EP300*, *FAT3,* and *CARD11* (Table S2)[45]. To our surprise, the mutation frequencies of *TP53* and *APC* were notably higher in the High-Mg group (Figure 2I).

**Figure 2.**
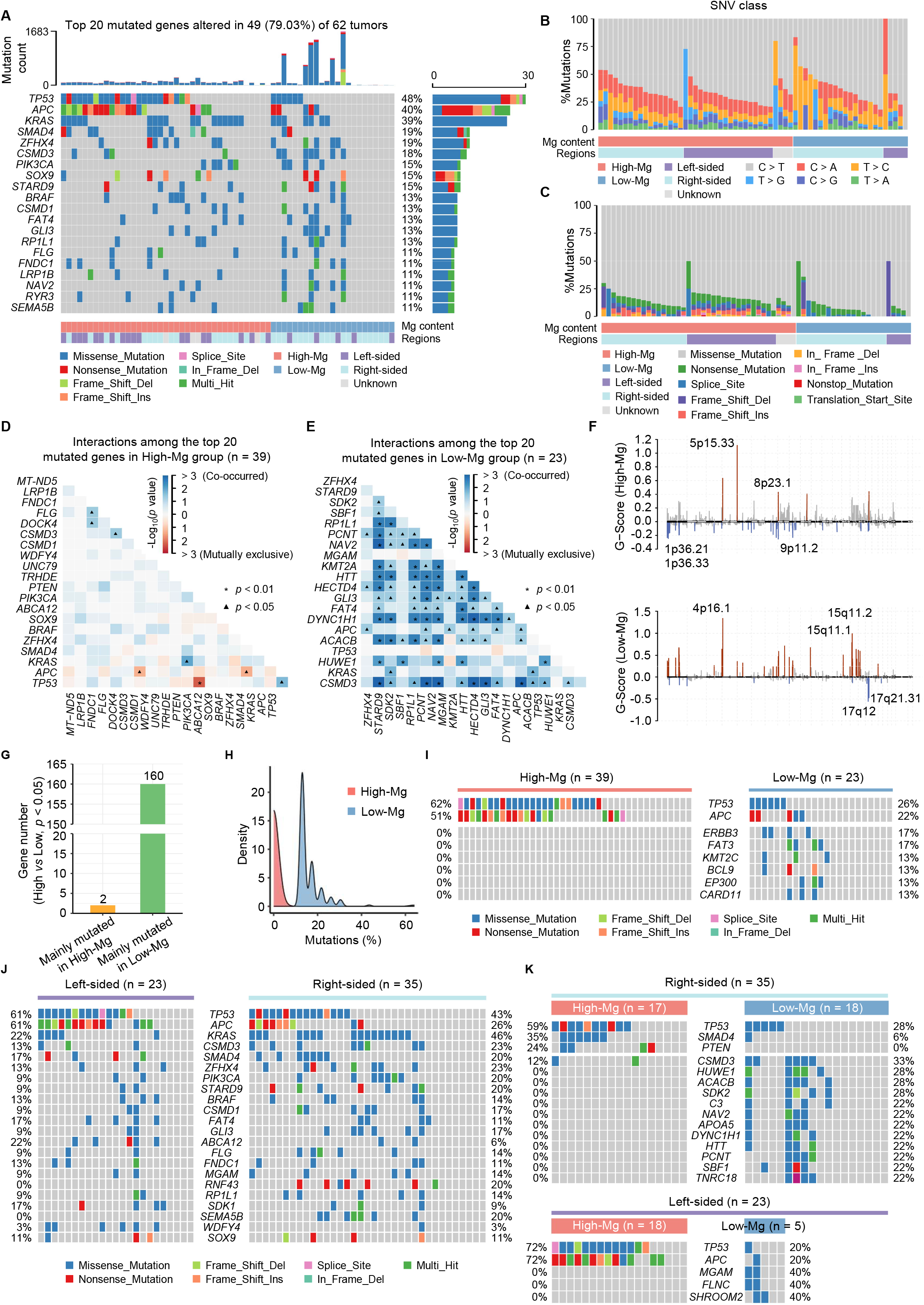
Low Mg content in tumors is linked to genome instability. **A.** Genetic profile of the top 20 mutated genes in High-Mg and Low-Mg tumors. Somatic mutations in the 20 genes were observed in 49 of 62 tumors. The top bar plot illustrates the overall count of somatic mutations in each patient, while the right bar plot depicts the distribution and composition of mutation types in each gene. **B.** Percentages of SNVs in High-Mg and Low-Mg groups. **C.** Percentages of the classes of mutations in High-Mg and Low-Mg groups. **D.** and **E.** The interactions among the top 20 mutated genes in the High-Mg (D) and Low-Mg (E) groups. *, *P* < 0.01; ▲, *P* < 0.05. Fisher’s exact test. **F.** Focal peaks exhibiting significant somatic copy-number amplification (red) and deletion (blue) (GISTIC2 Q-values < 0.1) are displayed in both High-Mg and Low-Mg groups. The top 5 amplified and deleted cytobands are labeled. **G.** Statistics of genes with significant variations in mutation frequencies between the High-Mg and Low-Mg groups. *P* < 0.05. Fisher’s exact test. **H.** Density plot displaying the mutation frequencies of genes in the High-Mg and Low-Mg groups. **I.** Genetic profile of the CRC driver genes with significant mutation frequencies between the High-Mg and Low-Mg groups. *P* < 0.05. Fisher’s exact test. **J.** Comparisons of gene mutation frequencies between left– and right-sided tumors. Genes with mutations in more than 7 out of 62 patients are displayed. **K.** Comparisons of gene mutation frequencies between High-Mg and Low-Mg tumors on the left (lower) or right (upper) side. Genes with mutations in more than 5 out of 62 patients are displayed.

Next, after excluding 4 tumors with unknown locations, we analyzed the impact of low Mg content on the genomic landscape of left-sided and right-sided colon cancer using the remaining 58 cases. Initially, we investigated the frequency of gene mutations in colon cancer and observed that the mutation frequencies of *TP53* and *APC* were higher on the left side, whereas *KRAS* exhibited a higher mutation frequency on the right side (Figure 2J), aligning with previous findings[29]. Subsequently, we explored the influence of Mg levels on gene mutations in left-sided and right-sided colon cancer separately. The results revealed that the majority of genes displayed higher mutation frequencies in the Low-Mg group on both sides. Notably, *TP53* demonstrated a higher mutation frequency in the High-Mg group for both left– and right-sided colon cancer, whereas *APC* exhibited a higher mutation frequency exclusively in the High-Mg group of left-sided tumors (72% cases in High-Mg group *vs* 20% cases in Low-Mg group) but not right-sided tumors (5 cases in High-Mg group *vs* 4 cases in Low-Mg group) (Figure 2K). Collectively, these results indicate that low Mg content in tumors probably increases the mutations of genes associated with CRC progression rather than initiation.

### Low Mg content in tumors is associated with EMT activation

To further explore the potential roles of Mg in CRC, we conducted a screening of the Mg-associated proteome in tumors according to a previously described strategy[37]. The correlation analysis of the levels of 5322 proteins and Mg content in the tumors revealed 1347 proteins that were positively correlated with Mg content and 1252 proteins that were negatively correlated with Mg content (**Figure 3**A; Table S3). Pathway enrichment analysis revealed that the 1252 proteins negatively correlated with Mg primarily belonged to cell adhesion-related pathways. Conversely, the 1347 proteins positively correlated with Mg were found to be associated with mRNA processing and translation pathways (Figure S3A−D). Further GSEA using 5322 proteins revealed that pathways negatively correlated with Mg content mainly included immune– and metastasis-related pathways (Figure 3B−D), such as the coagulation cascade, complement system, IL6-JAK-STAT3 signaling, TNFα signaling *via* NF-κB, EMT and angiogenesis (Figure 3B and D). On the other hand, pathways positively correlated with Mg content were predominantly related to cell proliferation and the cell cycle (Figure 3C).

**Figure 3.**
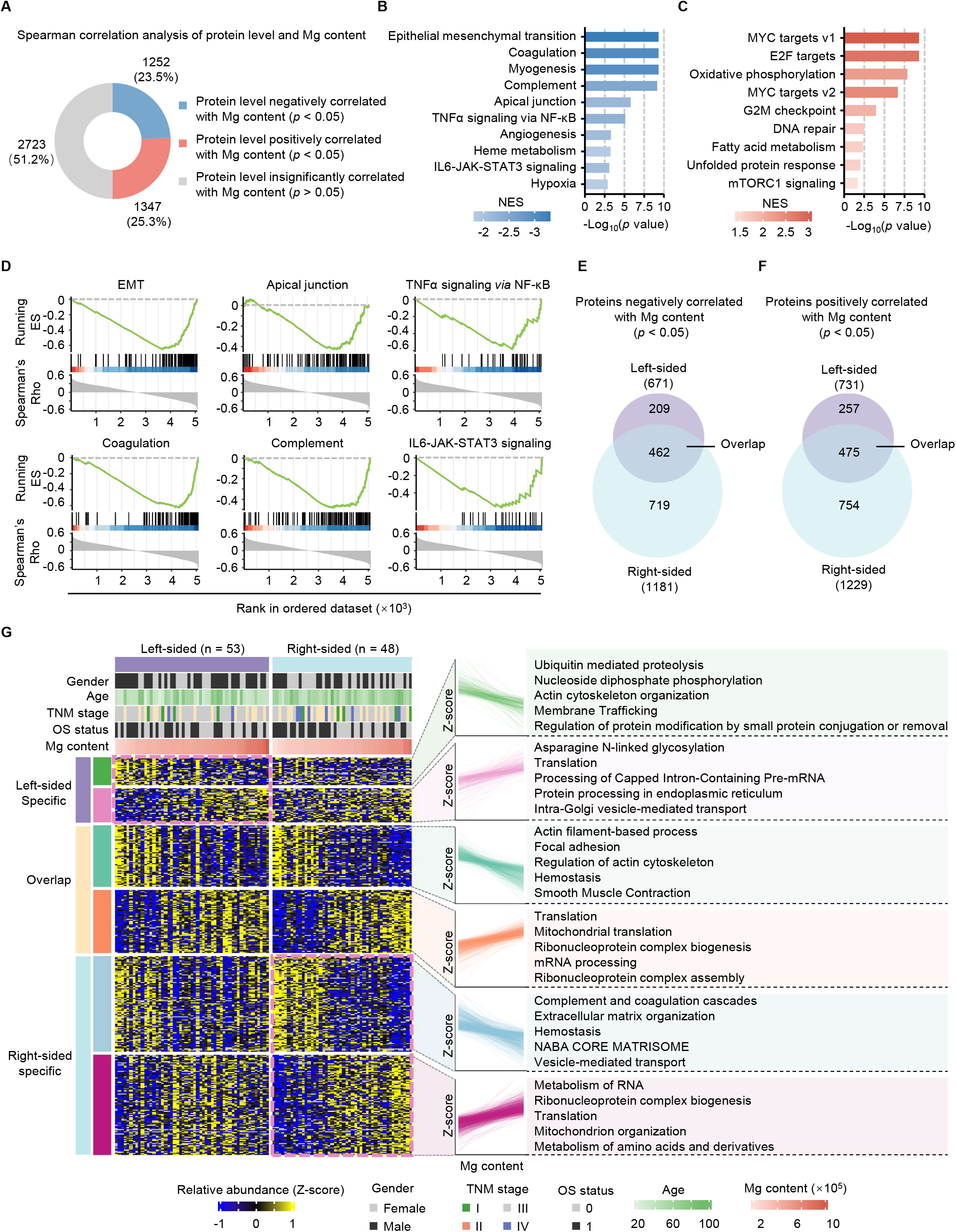
Low Mg content in tumors is associated with EMT activation. **A.** Spearman’s rank correlation analysis of protein levels and Mg content. GSEA pre-ranked hallmark analysis of the 5322 proteins. *P* < 0.05. Spearman’s rank correlation test. **B.** and **C.** Blue and red represent the pathways negatively (B) or positively (C) associated with Mg content, respectively. NES represents normalized enrichment score. **D.** Hallmark pathways related to immunity and metastasis. ES means enrichment score. **E.** and **F.** Venn diagrams displaying the overlapping proteins negatively (E) or positively (F) associated with Mg content between the left– and right-sided tumors. Spearman’s rank correlation analysis, *P* < 0.05. **G.** Heatmap showing the levels of Mg-related proteins in the left– and right-sided tumors. The correlation patterns of the proteins with Mg content in distinct modules are shown. Z-score of protein levels were mapped along Mg content in the left– and right-sided tumors. The top 5 pathways enriched by Metascape database using Mg-related proteins are shown. The gender, age, TNM stage, OS status and Mg content are annotated above the heatmap.

Additionally, we conducted separate screenings of Mg-associated proteins in left-sided and right-sided CRC (Figure 3E and F), and performed pathway enrichment analysis on these proteins (Figure 3G). The results indicated that proteins negatively correlated with Mg levels in either left-sided or right-sided CRC, or in both sides, were primarily associated with cell adhesion-associated pathways, such as regulation of actin cytoskeleton, actin cytoskeleton organization, focal adhesion, actin filament-based process, and extracellular matrix organization (Figure 3G). It is noteworthy that the complement and coagulation cascades were enriched exclusively in right-sided colon cancer, suggesting a more prominent influence of Mg on the inflammatory response in the right side (Figure 3G). As Mg have demonstrated variations between left-sided and right-sided CRC, we next asked whether Mg has impacts on different subtypes of CRC patients. To achieve this goal, we classified these patients into three subtypes utilizing the top 25% most variable proteins (Figure S4A−C). While the OS did not exhibit a significant difference among the three subtypes, further analysis indicated that subtype II displayed a comparatively lower OS rate and notably lower Mg content in comparison to subtypes I and III (Figure S4D and E). Proteins upregulated in subtype II (subtype II vs non-subtype II) were predominantly enriched in migration-related pathways (Figure S4F). Consistent with prior findings, Mg content closely correlates with the cell adhesion of CRC tumors.

Considering the findings from GSEA (Figure 3B and C), it can be inferred that low Mg content in tumors potentially affects tumor metastasis through the regulation of cell adhesion-related pathways. Consistent with our assumption, many proteins linked to cell-cell adhesion, such as tight junction proteins CLDN3, TJP2, and CGN, adherens junction proteins CDH1, CTNNB1, and CTNND1, and desmosome proteins DSC2, DSG2, and JUP, showed a significant positive correlation with Mg content. These proteins were crucial to maintain of the epithelial phenotype (**Figure 4**A; Table S4). In contrast, many cell-matrix adhesion proteins that were critical for maintaining the mesenchymal phenotype exhibited a significant negative correlation with Mg content, such as VCL, ITGA1, ITGB3, FN1, and FGA (Figure 4A). Further immunoblots confirmed increased N-cadherin, vimentin, vinculin, MMP2, and reduced E-cadherin, cingulin in Low-Mg tumors (Figure 4B), indicating the possible role of Mg in the EMT process.

**Figure 4.**
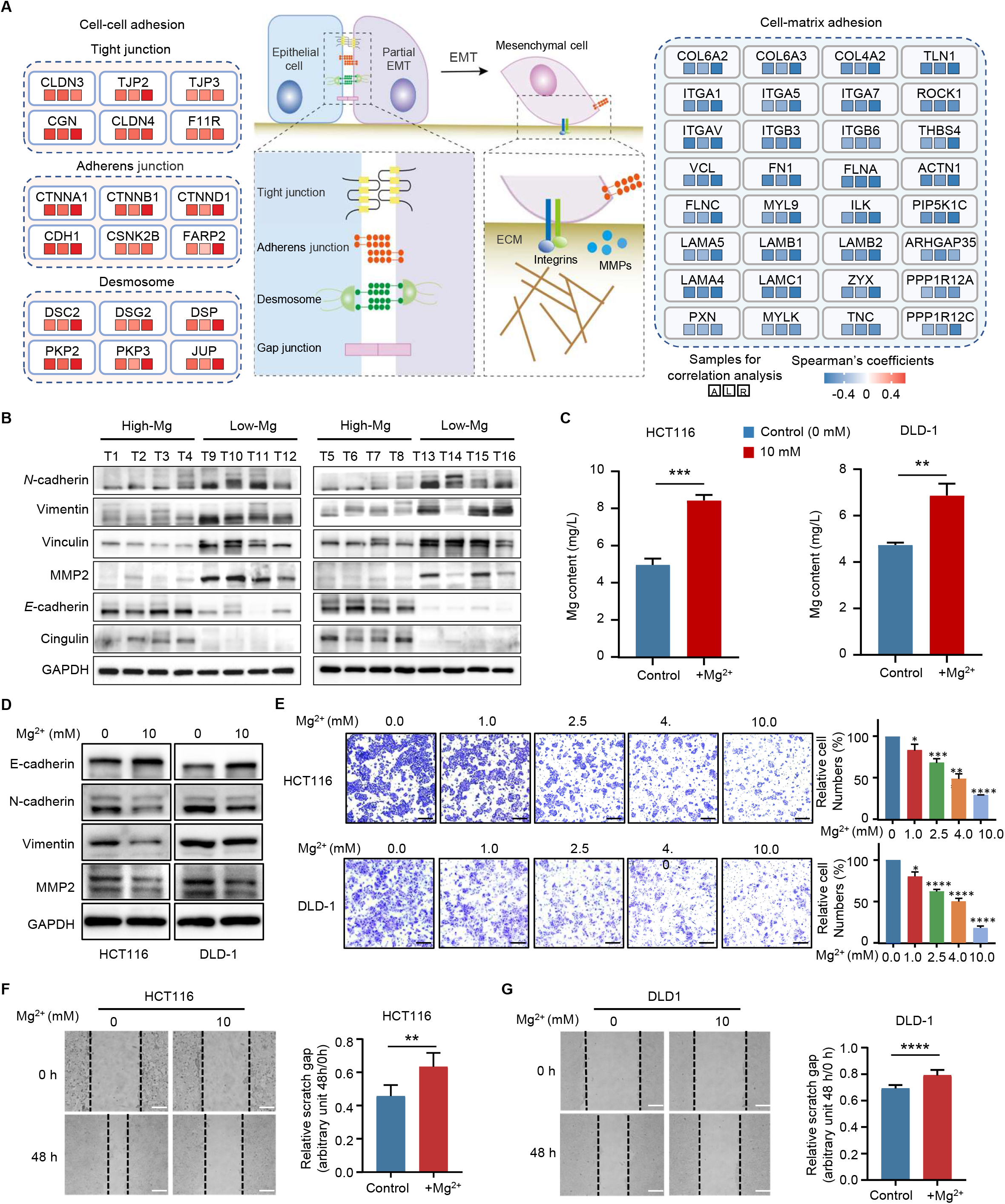
Functional validation of Mg in EMT activation in tumor cells. **A.** The diagram illustrates the schematic of the EMT and cell adhesion pathway. Spearman’s coefficients between Mg content and the levels of core components involved in cell-cell adhesion and cell-matrix adhesion are shown. The correlations were analyzed using all 104 tumor samples (A), 54 left-sided tumor samples (L), and 48 right-sided tumor samples (R). **B.** Immunoblots showing the expression of *N*-cadherin, vimentin, vinculin, MMP2, *E*-cadherin and cingulin in High-Mg and Low-Mg tumors. **C.** The measurements of Mg content in HCT116 and DLD-1 cells with or without MgCl_2_ treatment by ICP-MS. ***, *P* < 0.001; **, *P* < 0.01. Student’s *t* test. **D.** Western blotting analysis of vimentin, E-cadherin, N-cadherin and MMP2 in colon cancer cell lines with (10 mM) or without MgCl_2_ treatment. **E.** Representative images (left) and quantification results (right) of the migration assays using colon cancer cells treated with increasing concentrations of MgCl_2_. Scale bars, 500 μm. ****, *P* < 0.0001; ***, *P* < 0.001; **, *P* < 0.01; *, *P* < 0.05. Student’s *t* test. **F.** and **G.** Representative images (left) and quantification results (right) of the wound healing assays using HCT116 (F) and DLD-1 (G) cells treated with (10 mM) or without MgCl_2_. Scale bars, 200 μm. ****, *P* < 0.0001; **, *P* < 0.01. Student’s *t* test.

To validate the role of Mg in the regulation of tumor metastasis, transwell and wound healing assays were conducted using HCT116 and DLD-1 cell lines. ICP-MS analysis confirmed the successful uptake of Mg^2+^ into colon cancer cell lines (Figure 4C). Immunoblot analysis revealed that treatment with magnesium chloride (MgCl_2_) increased E-cadherin, and reduced the vimentin, MMP2, and N-cadherin expression in HCT116 and DLD-1 cells (Figure 4D). In line with the changes observed in EMT markers, both the transwell and wound healing assays demonstrated that treatment with MgCl_2_ significantly decreased the migratory ability of colon cancer cells (Figure 4E−G). Taken together, our findings suggest that low Mg content in tumors may activate EMT by disturbing the homeostasis of cell adhesion molecules.

### Potential mechanisms of EMT-related cell adhesion molecule alteration induced by low Mg content

Numerous studies have provided evidence indicating that inflammation is a key factor contributing to the loss of cell adhesion molecules[46−48]. Our proteomics analysis of Mg-associated proteins indicated that the expression of 44 complement and coagulation components, such as C1R, C1Q, C2, C3, and C5, was negatively correlated with Mg content and significantly increased in Low-Mg tumors and DNTs (Figure S3C)[49], suggesting the activation of the complement pathway in these tumors and paired DNTs[50], which is known to induce inflammation[51] and affect the expression of cell adhesion molecules (Figure S3D)[52−54].

Additionally, previous studies have shown that changes in protein phosphorylation are also important for the regulation of cell adhesion molecules[55, 56]. Mg^2+^ is an important regulator of protein phosphorylation. When Mg is deficient, it not only reduces intracellular ATP[30], but also impacts kinase and phosphatase activities[31],[32], leading to alterations in phosphorylation signaling in cancer cells. Therefore, in addition to its impact on maintaining the homeostasis of cell adhesion molecules by regulating inflammation, it is conceivable that low Mg content may also modulate the levels of cell adhesion molecules through the modulation of phosphorylation in cancer cells. To investigate the impact of Mg on tumors through phosphorylation mechanisms, we analyzed the Mg-associated phosphoproteome of colon cancer, leading to the identification of 332 and 465 phosphosites that were positively and negatively correlated with the Mg content in tumors, respectively (**Figure 5**A; Table S5). Enrichment analysis showed that proteins, such as ACTN4, DBN1, and VIM, corresponding to the 465 negatively correlated phosphosites were implicated in actin cytoskeleton organization, signaling by Rho GTPases, regulation of actin filament-based process, focal adhesion and cell-matrix adhesion (Figure 5B), while proteins, such as CDK11B, CDK12, and THRAP3, related to the 332 positively correlated phosphosites exhibited significant enrichment in tight junction and signaling by Rho GTPases pathways (Figure 5C), indicating that the Mg-associated phosphoproteome in colon cancer is closely linked to EMT-associated pathways (Figure 5D).

**Figure 5.**
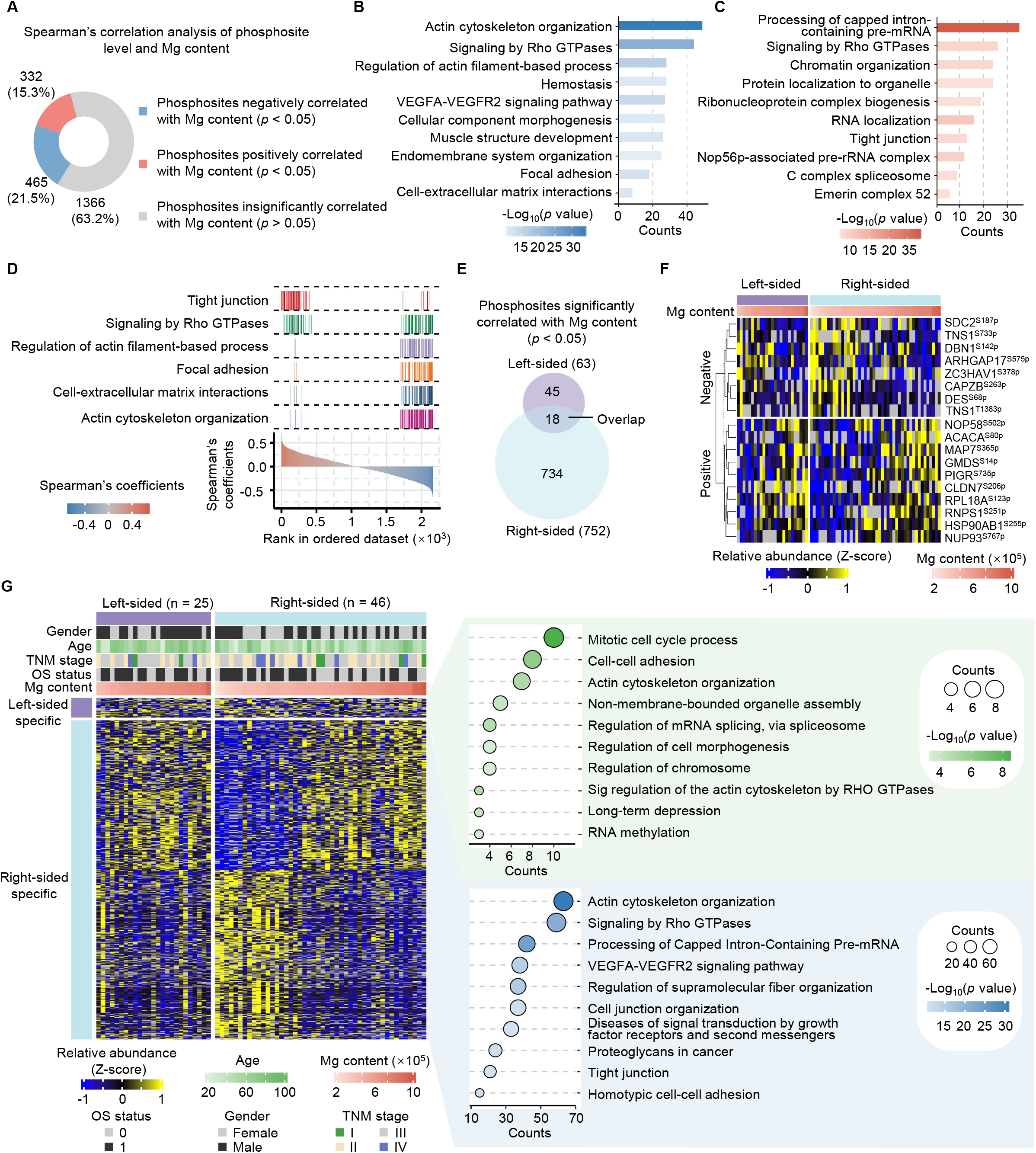
Impacts of Mg on the phosphoproteome of CRCs. **A.** Spearman’s rank correlation analysis of the levels of phosphosites and Mg content. *P* < 0.05. Spearman’s rank correlation test. **B.** and **C.** Pathway enrichment analysis of the proteins containing Mg negatively associated phosphosites (B) and positively associated phosphosites (C) by Metascape. **D.** Mg-correlated pathways related to cell adhesion are shown with Spearman’s coefficients obtained from (A). **E.** Venn diagrams displaying the overlapping phosphosites significantly correlated with Mg content between the left– and right-sided tumors. Spearman’s rank correlation analysis, *P* < 0.05. **F.** Heatmap displaying the levels of 18 common Mg-related phosphosites in left– and right-sided tumors. **G.** Heatmap showing the levels of phosphosites specifically correlated with Mg content in tumors on the left and right sides. The top 5 enriched pathways by Metascape using the proteins with Mg significantly correlated phosphosites are shown. Annotations above the heatmap include information such as gender, age, TNM stage, OS status, and Mg content. The heatmap illustrates the relative expression of phosphosites, utilizing the z-score for representation.

Furthermore, we separately screened Mg-associated phosphosites in left-sided and right-sided colon cancer. The results revealed that 18 phosphosites were significantly correlated with Mg content in both left-sided and right-sided colon cancer, while 45 phosphosites were only associated with Mg content in the left-sided colon cancer, and 734 phosphosites were specifically related to Mg content in the right-sided colon cancer (Figure 5E and F). Pathway enrichment analysis of the corresponding phosphorylated proteins unique to the left-sided and right-sided colon cancer demonstrated that the Mg-associated phosphorylated proteins on both sides were associated with cell adhesion pathways, including cell-cell adhesion, tight junction, cell junction organization, and actin cytoskeleton organization pathways (Figure 5G). Furthermore, through an integrative analysis of both the proteome and phosphorylome data, we identified 128 phosphosites positively correlated and 118 phosphosites negatively correlated with Mg content. These correlations were observed independently of protein levels (Figure S5A−C). Furthermore, enrichment analysis of phosphoproteins exhibiting significant correlations with Mg content highlighted their association with cell adhesion pathways, including actin cytoskeleton organization and cytoskeleton organization pathways (Figure S5D and E).

To validate the direct impacts of Mg on cancer cells via phosphorylation, we analyzed the phosphoproteome of colon cancer cells treated with and without MgCl_2_. As a result, of the 15,235 identified phosphosites, 564 sites were increased in the MgCl_2_-treated group, and 516 sites were reduced (**Figure 6**A and Table S6). Notably, cell adhesion pathways such as actin filament-based processes, regulation of actin cytoskeleton organization and signaling by Rho GTPases were enriched using proteins with differentially changed phosphosites (Figure 6B and C), consistent with the above phosphoproteomic data in CRC tissues (Figure 5B and C). Overlapping analysis indicated that the expression of 28 phosphosites were significantly changed with Mg content in both HCT116 and CRC tissues (Figure 6D and E). Among them, 4 phosphosites, including DBN1^S142p^, ARHGAP1^S51p^, TNS1^S1177p^, and FLNA^S1459^, were involved in cell adhesion (Figure 6E). Additionally, a high degree of phosphorylation at these four sites was found to be a predictor of unfavorable prognosis among CRC patients (Figure 6E and F, Figure S6A). Taken together, our results indicate that low Mg content in tumors can reshape the protein phosphorylation network of proteins associated with EMT and may ultimately affect tumor cell migration.

**Figure 6.**
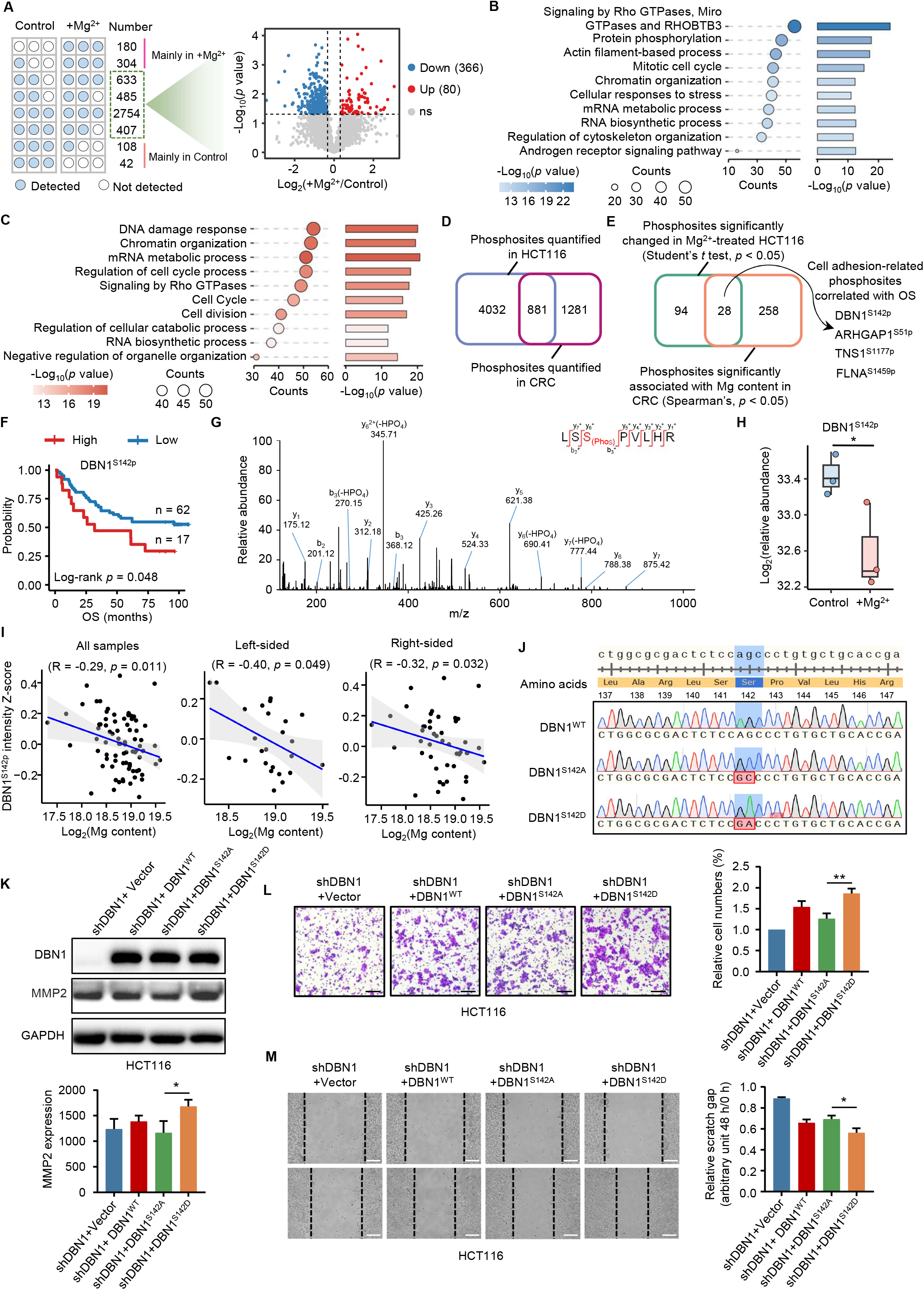
Mg-regulated DBN1^S142p^ reduction inhibits cell migration in CRC cells. **A.** The statistics of the number of phosphosites occurring in the +Mg^2+^ group or in the control group (left). The phosphosites mainly identified in the Mg^2+^ and control groups were directly considered upregulated (484) and downregulated (150), respectively. Additionally, Volcano plot showing 80 upregulated and 366 downregulated phosphosites in the +Mg^2+^ group and control group (right), respectively, using phosphosites identified in at least two replicates of each group. *P* < 0.05. Cutoff, ratio(+Mg^2+^/Control) > 1.2 or < 0.83. Student’s *t* test. Blue and red dots represent significantly increased phosphosites in the control and +Mg^2+^ groups, respectively. **B.** and **C.** Pathway enrichment analysis of the corresponding proteins of downregulated (516) (B) and upregulated (564) (C) phosphosites in the +Mg^2+^ group using Metascape. **D.** Venn diagram showing the number of overlapping phosphosites quantified in HCT116 and CRC. **E.** Venn diagram showing the number of overlapping phosphosites with significant changes in Mg^2+^-treated HCT116 cells and significant correlations with Mg content in CRC. **F.** Survival analysis of 79 CRC patients with different DBN1^S142p^ levels in tumors. *P* < 0.05. Log-rank test. **G.** MS/MS spectrum of the identified phosphopeptide containing DBN1^S142p^. **H.** The statistics of the levels of DBN1^S142p^ in the control and +Mg^2+^ groups. *, *P* < 0.05. Student’s *t* test. **I.** Spearman’s rank correlation analysis of DBN1^S142p^ and Mg content in all (72), left (25)– and right (46)-sided tumors. **J.** Mutation information of the DBN1^WT^, DBN1^S142A^ and DBN1^S142D^ genes and sequencing results. **K.** Immunoblotting assays determined the effects of DBN1 mutations on MMP2 expression. *, *P* < 0.05. Student’s *t* test. **L.** and **M.** Transwell migration (L) and wound healing assays (M) were performed in cells transfected with vector, DBN1^WT^, DBN1^S142A^ and DBN1^S142D^. Scale bars, 500 μm (L) and 200 μm (M). **, *P* < 0.01; *, *P* < 0.05. Student’s *t* test.

### Mg-regulated DBN1^S142p^ reduces the interaction between DBN1 and ACTN4 and contributes to EMT

DBN1 is a protein that binds to F-actin, which is essential for maintaining the levels of cell adhesion molecules[57, 58]. However, the functions of DBN1^S142p^ in CRC and the mechanism by which Mg-regulated DBN1^S142p^ contributes to the EMT process are completely unknown. The confident identification of DBN1^S142p^ was verified through the MS/MS spectrum (Figure 6G). Statistical analysis revealed a significant decrease in DBN1^S142p^ in the MgCl_2_-treated group (Figure 6H). Moreover, a remarkable negative correlation was observed between DBN1^S142p^ and Mg content in CRC, irrespective of its sidedness (Figure 6I). To explore the role of DBN1^S142p^ in the regulation of EMT, we transfected wild-type DBN1 or DBN1 mutants in *DBN1*-knockdown HCT116 cells that mimic the phosphorylation (S142D) or de-phosphorylation (S142A) states (Figure 6J). Given that DBN1 is an F-actin-binding protein and that the assembly of F-actin can promote MMP2 expression[57, 58], we assumed that the Mg-regulated decrease in DBN1^S142p^ might serve as a signal triggering F-actin disassembly and promoting MMP2 degradation. Immunoblotting results showed that DBN1^S142D^ remarkably upregulated MMP2 compared with DBN1^S142A^ (Figure 6K). Consistently, both transwell and wound healing assays demonstrated that the DBN1^S142A^ mutation significantly inhibited cell migration (Figure 6L and M). Further studies showed that MgCl_2_ treatment, which was able to reduce the level of DBN1^S142p^ (Figure 6H), obviously enhanced the formation of F-actin (**Figure 7**A, Figure S6B). Consistent with these findings, DBN1^S142A^, which mimics the de-phosphorylation state, increased the formation of F-actin compared with DBN1^S142D^ (Figure 7B). In addition, we further explored the function of ARHGAP1^S51A^. While the ARHGAP1^S51A^ mutation also notably inhibited cell migration, it did not impact F-actin formation in contrast to ARHGAP1^S51D^. This suggests that ARHGAP1^S51p^ inhibits cell migration through mechanisms other than F-actin formation (Figure S6C−E).

**Figure 7.**
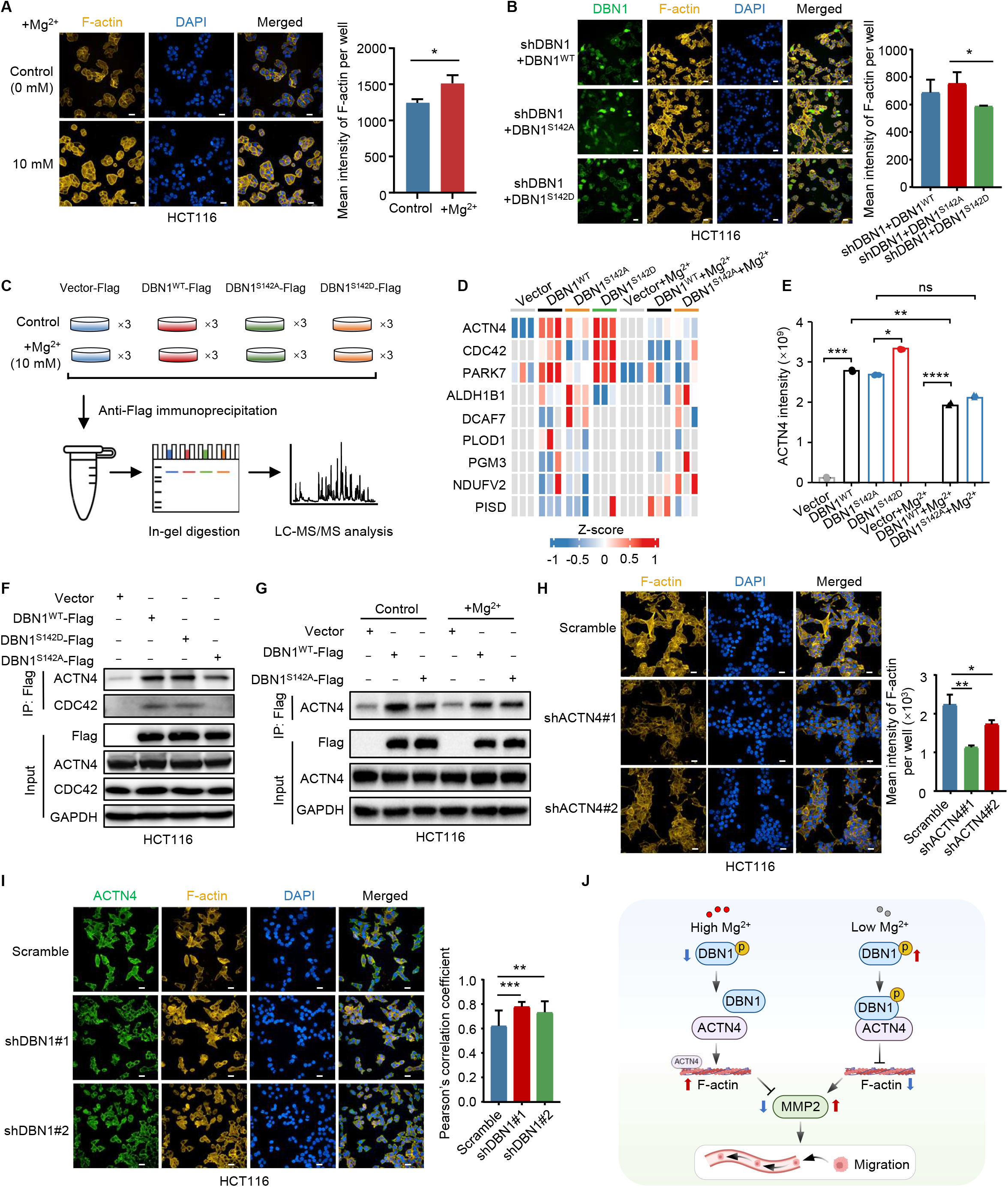
Mg-regulated DBN1^S142p^ reduces the interaction between DBN1 and ACTN4 and contributes to EMT. **A.** Effects of MgCl_2_ treatment on the formation of F-actin determined by immunofluorescence assays in HCT116 cells. Scale bars, 500 μm. *, *P* < 0.05. Student’s *t* test. **B.** Effects of DBN1^WT^, DBN1^S142A^ and DBN1^S142D^ on the formation of F-actin determined by immunofluorescence assays. Scale bars, 500 μm. *, *P* < 0.05. Student’s *t* test. **C.** Schematic representation of the identification of DBN1-binding proteins affected by DBN1^S142p^ and Mg^2+^ through affinity purification followed by MS analysis. **D.** Heatmap showing 9 DBN1-binding proteins affected by Mg^2+^ through regulating DBN1^S142p^. Z-score of protein levels were used. **E.** The relative intensity of the DBN1-binding protein ACTN4 determined by MS in immunoprecipitation assays under different experimental conditions. ****, *P* < 0.0001; ***, *P* < 0.001; **, *P* < 0.01; *, *P* < 0.05; ns, means no significance. Student’s *t* test. **F.** Immunoblotting assays determined the effects of DBN1^S142p^ on the interactions of ACTN4 and CDC42 with DBN1. **G.** Immunoblotting assays determined the effects of Mg on the interaction between ACTN4 and DBN1 affected by DBN1^S142p^. **H.** Effects of ACTN4 on the formation of F-actin determined by immunofluorescence assays. Scale bars, 500 μm. **, *P* < 0.01;*, *P* < 0.05. Student’s *t* test. **I.** Co-localization of ACTN4 and F-actin was determined by immunofluorescence assays. Scale bars, 500 μm. ***, *P* < 0.001; **, *P* < 0.01. Student’s *t* test. **J.** Potential mechanism of Mg-regulated cell migration *via* modulation of DBN1^S142p^.

To elucidate the impact of Mg-regulated DBN1^S142p^ on F-actin polymerization, we employed immunoprecipitation coupled MS analysis. This approach allowed us to identify proteins binding to DBN1, the interactions of which were influenced by DBN1^S142p^ and Mg^2+^ (Figure 7C). As a result, we identified 376 proteins that bind to DBN1, with 30 and 46 proteins being enriched in the DBN1^S142D^ and DBN1^S142A^ groups, respectively (Figure S7A and B; Table S7). Notably, some of these proteins are involved in actin assembly and disassembly, suggesting that the level of DBN1^S142p^ may affect its binding to cytoskeleton-related proteins. Upon comparing the differential binding proteins between the DBN1^WT^ and DBN1^WT^+Mg^2+^ groups, we noted that 27 proteins exhibited increased interactions with DBN1, while interactions with 56 proteins were diminished in response to MgCl_2_ treatment (Figure S7C). Of the 83 differential proteins, 73 proteins demonstrated no notable changes between the DBN1^S142A^ and DBN1^S142A^+Mg^2+^ groups (Figure S7C), indicating that Mg^2+^ might modulate the interactions of these 73 proteins with DBN1 by altering DBN1^S142p^. Among these 73 proteins, 53 whose interactions were weakened by Mg^2+^ were primarily associated with the actin cytoskeleton (Figure S7D), indicating that Mg^2+^ could affect the interaction between DBN1 and these cytoskeletal proteins, which may further impact the depolymerization of F-actin. Additional comparisons were performed to identify the proteins that bind to DBN1 in both the phosphorylated and dephosphorylated states at the S142 residue. (Figure S7E and F). The results indicated that Mg suppressed DBN1^S142p^ to inhibit the interaction of three proteins, namely, ACTN4, CDC42, and PARK7 (Figure 7D and E, Figure S7E), while it enhanced the interaction of six proteins, namely, ALDH1B1, DCAF7, PLOD1, PGM3, NDUFV2, and PISD (Figure 7D, Figure S7 F and G). Among these proteins, ACTN4 and CDC42 are well known for their close association with the cell cytoskeleton[59].

Previous studies have indicated that ACTN4 participates in the formation of F-actin and that F-actin stability is compromised when ACTN4 is detached from F-actin[59]. Thus, we propose that the interaction of DBN1 to ACTN4 results in the detachment of ACTN4 from F-actin. The de-phosphorylation of DBN1 at S142 could weaken the interaction between DBN1 and ACTN4, thereby promoting the interaction of ACTN4 to F-actin and enhancing F-actin stability. To investigate this assumption, we overexpressed Flag-tagged vector, DBN1^WT^, DBN1^S142A^, and DBN1^S142D^ in *DBN1*-knockdown HCT116 cells and performed immunoprecipitation under identical experimental conditions. Immunoblotting results confirmed ACTN4’s ability to interact with DBN1 (Figure 7E). Moreover, the interaction between the mutant DBN1^S142A^ and ACTN4 was weakened, indicating that DBN1^S142p^ may affect the DBN1-ACTN4 interaction (Figure 7F).

To confirm that Mg^2+^ reduces the interaction between DBN1 and ACTN4 by decreasing the level of DBN1^S142p^, we constructed stable cell lines overexpressing Flag-tagged vector, DBN1^WT^, and DBN1^S142A^. These cell lines were treated with and without MgCl_2_, followed by Flag-targeted immunoprecipitation assays. The immunoblotting results showed that MgCl_2_ treatment weakened the interaction between DBN1^WT^ and ACTN4, while DBN1^S142A^-ACTN4 remained consistent after MgCl_2_ treatment (Figure 7G). These findings imply that Mg^2+^ may decrease the interaction between DBN1 and ACTN4 by reducing DBN1^S142p^. Then, we generated stable HCT116 cell lines with *ACTN4* knockdown and examined the effect of ACTN4 on F-actin polymerization (Figure S7H). The results showed that *ACTN4* knockdown inhibited F-actin formation (Figure 7H). In contrast, knockdown of *DBN1* resulted in a significant increase in colocalization between ACTN4 and F-actin (Figure 7I), suggesting that the interaction between DBN1 and ACTN4 inhibits the formation of F-actin.

Collectively, these findings suggest that Mg^2+^ diminishes the interaction between DBN1 and ACTN4 by decreasing the level of DBN1^S142p^. This, in turn, enhances the binding of ACTN4 to F-actin and stabilizes F-actin, ultimately resulting in reduced MMP2 expression and decreased migratory ability of colon cancer cells (Figure 7J).

## Discussion

In summary, for the first time, we presented an atlas of the Mg-associated genome, proteome and phosphoproteome in CRC and demonstrated that low Mg content in tumors promoted genome instability and tumor metastasis. This work has yielded some novel findings. First, our study revealed that a large number of genes in Low-Mg tumors exhibited higher mutation frequencies and co-occurred, including several driver genes associated with tumor progression. Conversely, High-Mg tumors displayed high mutation frequencies in well-known initiation driver genes of CRC, such as *TP53* and *APC*, indicating that the lack of Mg content may primarily impact tumor progression rather than initiation at the genomic level. Second, our study showed that Low-Mg tumors activated EMT by disrupting the homeostasis of adhesion molecules in cancer cells. This disruption was attributed not only to complement pathway activation and elevated inflammation in Low-Mg tumors but also to changes in the phosphorylation signaling of proteins associated with cell adhesion. Third, we discovered a novel phosphosite, DBN1^S142p^, that is regulated by Mg^2+^. Our results showed that Mg^2+^ reduced the interaction between DBN1 and ACTN4 by decreasing DBN1^S142p^. This reduction in DBN1^S142p^ resulted in increased binding of ACTN4 to F-actin, thereby stabilizing F-actin and ultimately leading to reduced MMP2 expression and inhibition of colon cancer cell migration. Overall, this work uncovers the roles of Mg in CRC and demonstrates that low Mg content in tumors could be a potential driver factor of CRC.

Cancer cells require a large amount of Mg to support their glycolytic metabolism, ATP production and protein synthesis, which are essential for sustaining rapid proliferation[60]. Insufficient Mg uptake not only slows cancer cell proliferation but also leads to genome instability[61], as we demonstrated in this work. Mg deficiency is known to induce low-grade systemic inflammation and cause increased production of ROS and proinflammatory cytokines such as IL-6, TNF-α, and IL-1β[8, 62], leading to genome instability. Moreover, Mg also plays a crucial role in regulating DNA replication and repair processes[39, 40, 42−44], which may partially explain why a significant number of CRC driver genes, such as *KMT2C* and *ERBB3*, are mutated in tumors with low Mg content. Tumor initiation and progression are determined by the inactivation of the tumor suppressors *APC* and *TP53* and the activation of the oncogene *KRAS*[63]. Surprisingly, the mutation frequencies of these genes were much higher in tumors with high Mg content (High-Mg/Low-Mg frequencies: *APC*, 51%/22%, *P* = 0.032; *TP53*, 62%/26%, *P* = 0.0091; *KRAS*, 44%/30%, *P* = 0.42). However, the mechanism by which high levels of Mg induce mutations in the *APC* and *TP53* genes remains to be investigated. In addition, the screening of Mg-related gene mutations in left– or right-sided colon cancer, or on both sides, requires validation in larger CRC cohorts.

EMT, characterized by loss of cell-cell junctions, disruption of cell-matrix attachments and cytoskeleton remodeling, can increase the mobility of cells. Many signaling pathways, such as TGFβ, Wnt and Hippo, are associated with the regulation of EMT[64]. Mg deficiency is well known to cause inflammation by disturbing the coordination of the innate immune system and the adaptive immune response[14], leading to rapid loss of cell adhesion molecules[46−48]. Alternatively, phospho-signals, serving as the first wave of response to intracellular and extracellular changes, extensively exist during the EMT process[65, 66]. For example, phosphorylation of SNAIL[67], GSK3β[68], and EGFR[69] contributes to the regulation of EMT. Our findings, for the first time, systematically highlight the significance of Mg’s direct regulation of phosphorylation in cancer cells, which is critical to maintain the homeostasis of cell adhesion molecules and is closely linked to the EMT process. Specifically, our experiments revealed that Mg^2+^ played a crucial role in reducing the interaction between DBN1 and ACTN4 by lowering the level of DBN1^S142p^. This, in turn, had a significant impact on F-actin stability and the homeostasis of cell adhesion molecules. However, the exact mechanism by which Mg affects DBN1^S142p^ is not yet fully understood, although it may impact the activity of phosphatases, thereby influencing DBN1^S142p^. Additionally, the precise binding mode between DBN1 and ACNT4 remains elusive, and further biochemical experiments are necessary to determine how DBN1^S142p^ affects the interaction between DBN1 and ACNT4.

Adults typically consume a daily Mg intake of approximately 330−350 mg. Previous studies have showed that cirrhotic patients with hepatocellular carcinoma (HCC) exhibit reduced serum Mg levels compared to those without HCC, independent of confounding factors such as dietary Mg intake and medications affecting Mg levels[70]. Moreover, mouse experiments have validated that the growth of primary tumors sequesters Mg from the extracellular environment, leading to hypomagnesemia[60]. Additionally, both our work and other studies have described the higher preference of Mg absorption in tumors than normal tissues, which may raise the possibility of reducing serum Mg levels[60, 70]. However, this speculation requires further evidence.

Previous clinical studies have demonstrated the potential benefits of Mg in the prevention and treatment of CRC. A population-based prospective study suggests that a high Mg intake may reduce the risk of CRC in women[21]. Moreover, Mg supplementation can enhance the effect of drug treatment and minimize serious side effects in CRC patients. For example, a combination of 25(OH)D3 and Mg is essential in reducing the risk of mortality in CRC patients[71]. Peripheral neuropathy is a common side effect caused by chemotherapy in CRC patients, but a high dietary Mg intake can reduce its prevalence and severity[28]. Notably, in late-stage CRC patients treated with cetuximab[72] or bevacizumab[73], EGFR inhibition can result in Mg wasting due to decreased renal reabsorption, and circulating Mg^2+^ reduction may act as a predictive factor of treatment efficacy and outcome[74, 75]. The extracellular matrix (ECM) is a complex network of proteins and other molecules that surround cells and play important roles in cell signaling, migration, and proliferation[76]. Abnormalities in ECM composition and organization are often associated with cancer progression and metastasis. ECM proteins themselves and ECM protein-interacting proteins such as integrins are known targets of cancer treatment[77, 78]. Interestingly, by referring to known drug targets with FDA-approved drugs or candidate drugs in clinical trials, we identified 24 clinically actionable cell-matrix proteins, such as COL3A1, FGA, and ITGA5, whose expression was negatively correlated with Mg content in tumors (Spearman’s rank correlation test, *P* < 0.05, Figure S8; Table S8), suggesting that Mg might serve as an adjuvant drug for precise CRC treatment by affecting the levels of these potential drug targets. Our research revealing the functions of Mg is expected to promote the application of Mg reagents in the prevention and treatment of CRC.

## Materials and methods

### Sample collection and preparation

A total of 115 paired tumors and corresponding DNTs wereobtained from treatment-naive CRC patients at West China Hospital of Sichuan University. The samples were rapidly snap-frozen in liquid nitrogen and then stored at –80 °C for long-term preservation. This study was carried out with the approval of the Research Ethics Committee (Approval No. 2020 (374)). Informed consent and approvals were obtained from each patient and reviewed accordingly. Detailed clinical information, such as age, gender, tumor region, OS status, OS (months) and TNM stage, is systematically recorded in Table S1. Patient follow-up was conducted over a median period of 67.25 months. OS was the duration from surgery to either the patient’s demise or the last follow-up visit. We used genomic, proteomic, and phosphoproteomic data of these CRC patients to elucidate the impact of Mg deficiency on CRC. Notably, the genome, proteome, and phosphoproteome data for each patient were generated using a same sample.

### Proteomic and phosphoproteomic analyses

Protein extraction and digestion procedures were as follows: Tissues were homogenized and lysed using gentleMACS Dissociators (Catalog No. 130-093-235, Miltenyi Biotec, Nordrhein-Westfalen, Germany) with RIPA buffer (Catalog No. P0013C, Beyotime, Shanghai, China). Cell samples, on the other hand, were directly lysed in RIPA buffer. Subsequently, the prepared lysates were sonicated for 5 min at 227.5 W, with a 3-second on and 10-second off cycle. After centrifugation at 20,000 g for 20 min at 4 °C, the supernatant was transferred to a new tube, and the Bradford protein assay was taken to measure protein concentration. For each sample, 100 μg of protein lysates were first reduced using 10 mM tris (2-carboxyethyl) phosphine (TCEP) at 56 °C for 60 min. Subsequently, alkylated with 20 mM iodoacetamide for 30 min in the dark at 25 °C. The protein samples were subsequently precipitated using CH_3_OH, CHCl_3_, and H_2_O (CH_3_OH: CHCl_3_: H_2_O = 4:1:3, v/v). Finally, the proteins were digested with trypsin at a 1:50 (trypsin/protein, w/w) for 12 hours.

Tandem Mass Tag (TMT)-10 labeling of peptides was conducted as follows: The TMT-10 plex Isobaric Label Reagent (Catalog No. 90110, Thermo Fisher Scientific, Waltham, MA) was dissolved in anhydrous acetonitrile (ACN) after reaching room temperature. 10 μg and 40 μg digested peptides of each sample for proteome and phosphoproteome analysis were labeled with TMT reagents following the manufacturer’s instructions, respectively. Subsequently, the TMT-labeled peptides were combined and dried.

TMT-labeled peptide fractionation was performed as follows: Reversed-phase high-performance liquid chromatography (RP-HPLC) was used to fractionate peptides with a basic mobile phase in proteomics. The separation was carried out with a flow rate of 1 mL/min using a mixture of buffer A (98% H_2_O, 2% ACN, pH = 10) and buffer B (90% ACN, 10% H_2_O, pH = 10). The LC gradient run spanned 120 min and followed this pattern: 3%−35% buffer B in 95 min, 35%−60% buffer B in 10 min, 60%−100% buffer B in 15 min. The eluates were collected in 120 fractions, which were subsequently merged into 15 fractions for each batch of CRC samples. These fractions were then dried using a vacuum centrifuge. Following desalting with C18 ZipTips, the TMT-labeled peptides were subjected to LC-MS/MS analysis. For phosphoproteomics, the TMT-labeled peptides from tissue samples were initially fractionated into 15 fractions using a C18 solid-phase extraction (SPE) columns (100 mg/1 mL). Later, these fractions were further combined into 5 fractions and dried. In the case of cell phosphoproteomics, each sample was divided into 9 fractions using a C18 SPE columns, which were then merged into 3 fractions prior to drying.

The enrichment of phosphorylated peptides was performed as follows: Phosphopeptides were enriched using PureCube Fe-NTA Agarose Beads (Catalog No. 31403-Fe, Cube Biotech, Monheim, Germany) according to the manufacturer’s instructions. The peptides, dissolved in 300 µL of loading buffer (85% ACN, 0.1% TFA), were then incubated with the prepared beads for about 60 min on a 3D shaker at room temperature. Subsequently, the agarose beads were washed 4 times with washing buffer (80% ACN, 0.1% TFA) and then eluted with 150 µL of elution buffer (composed of 40% ACN and 15% ammonium hydroxide). To neutralize the eluate, 8 µL of 20% TFA was added. The elution buffer were subsequently dried by vacuum, and subjected to LC-MS/MS analysis after desalting using C18 ZipTips.

For proteomics, LC-MS/MS analysis was conducted using a Nano EASY-nLC 1200 liquid chromatography system LC instrument coupled with an Orbitrap Exploris 480 mass spectrometer (Catalog No. BRE725539, Thermo Fisher Scientific). After desalting with a Ziptip columns, the samples were dried and reconstituted in buffer A (98% H_2_O, 2% ACN, 0.1% formic acid (FA)). They were then loaded onto an in-house pulled and packed analytical column (75 μm × 30 cm) packed with C18 particles. Samples were analyzed using a 65-minute gradient of 4% to 100% buffer B (0.1% FA in 80% ACN) at a flow rate of 300 nL/min in positive ion mode. The MS1 full scans (m/z 350−1800) were acquired with a resolution of 60,000. The automatic gain control (AGC) value was set to 300%, and the maximum injection time (MIT) was 50 ms. For MS/MS scans, the top 20 most abundant parent ions were selected under an isolation window of 0.7 m/z, and fragmentation was performed using a normalized collision energy (NCE) of 36%. The normalized AGC value for MS/MS was set to 75%, and the MIT was 80 ms. Precursor ions with charge states of z = 1, 8, or unassigned charge states were excluded from further fragmentation.

For phosphoproteomics, LC-MS/MS analysis was conducted using a Nano EASY-nLC 1200 liquid chromatography system LC instrument coupled with a Q Exactive HF-X high-resolution mass spectrometer (Catalog No. BR64966, Thermo Fisher Scientific). After desalting with Ziptip columns, the samples were dried and reconstituted in buffer A, consisting of 2% ACN and 0.1% FA. They were then separated by a homemade trap column (2.5 cm × 75 μm) packed with Spursil C18 particles and an analytic column (25 cm × 75 μm) packed with Reprosil-Pur C18-AQ particles. Samples were analyzed using a 65-min gradient of 6% to 100% buffer B (0.1% FA in 80% ACN) at a flow rate of 330 nL/min in positive ion mode. The MS1 full scans (m/z 350−1600) were acquired with a resolution of 60,000. The AGC value was set to 3e6, and the MIT was 20 ms. For MS/MS scans, the top 20 most abundant parent ions were selected under an isolation window of 0.6 m/z, and fragmentation was performed using stepped NCE of 25% and 31%. Precursor ions with charge states of z = 1, 8, or unassigned charge states were excluded from further fragmentation. For label-free cell phosphoproteomics, samples were analyzed using a 90-min gradient of 12% to 100% buffer B (0.1% FA in 80% ACN) at a flow rate of 330 nL/min in positive ion mode. The MS1 full scans (m/z 350−1800) were acquired with a resolution of 60,000. The AGC value was set to 3e6, and the MIT was 20 ms. For MS/MS scans, the top 20 most abundant parent ions were selected under an isolation window of 1.6 m/z, and fragmentation was performed using stepped NCE of 25% and 27%. Precursor ions with charge states of z = 1, 8, or unassigned charge states were excluded from further fragmentation.

### MS database searching

All mass spectrometry raw data files of proteomics and phosphoproteomics were analyzed by using MaxQuant (version 1.6), aligned against the Swiss-Prot human protein sequence database comprising 20,413 entries (updated 04/2019). For MS2 reporter ion quantification, the reporter mass tolerance was set at 0.02 Da. Peptide mass tolerance was set at 10 ppm, and only peptides and proteins with a false discovery rate (FDR) lower than 1% were kept for further data processing. Up to 2 trypsin-missing cleavage sites were allowed. Cysteine carbamidomethylation was specified as a fixed modification, while oxidation of methionine and protein N-terminal acetylation were considered variable modifications. For phosphoproteomics analysis, phosphorylation (+79.9663 Da) of serine, threonine, and tyrosine residues were also added to the above variable modifications.

### Data cleaning of proteome data

R (version 4.2.1) was used to process proteome data to minimize systematic errors based on the “peptides.txt” from MaxQuant output. Several preprocessing steps were performed to refine the data. First, potential contaminants and reverse proteins were excluded. Subsequently, only proteins with ≥ 2 unique peptides were selected for further analysis. To ensure comparability across samples within the same batch, the total protein abundance of each sample was adjusted to an equal level. To reduce the impact of noise, the protein intensity values of the tumor or DNT samples were divided by the protein intensity of an IS sample, yielding protein sample-to-standard (S/S) values. This step helped normalize the data and mitigate potential confounding factors. All data from the 27 batches were combined into a matrix, with samples represented as columns and proteins as rows. Additionally, the normalized values underwent an additional log2-transformation to facilitate subsequent analyses. The values in the matrix were transformed to column z-scores and row z-scores to normalize the data distribution across samples and proteins, respectively. Any values that were originally zero were considered missing and replaced with “NA”. To ensure data quality, proteins with > 50% missing values were excluded from the dataset, resulting in a refined and reliable set of proteins for further analysis.

### Data cleaning of phosphoproteome data

R (version 4.2.1) was used to process phosphoproteome data to minimize systematic errors based on the “Phospho (STY) Sites.txt”. Several preprocessing steps were performed to refine the data. First, potential contaminants and reverse peptides were excluded. In addition, phosphopeptides with localization probability > 0.75 were kept. To ensure comparability across samples within the same batch, the total intensities of phosphopeptides in each sample was adjusted to an equal level. To reduce the impact of noise, the phosphopeptide intensities of the tumors or DNT samples were divided by the phosphopeptide intensity of an IS sample, resulting in sample-to-standard (S/S) values. This step helped normalize the data and mitigate potential confounding factors. All data from the 20 batches were combined into a matrix, with samples represented as columns and phosphopeptides as rows. Additionally, the normalized values underwent an additional log2-transformation to facilitate subsequent analyses. The values in the matrix were transformed to column z-scores and row z-scores to normalize the data distribution across samples and phosphopeptides, respectively. Any values that were originally zero were considered missing and replaced with “NA”. To ensure data quality, phosphopeptides with > 50% missing values were excluded from the dataset.

### Measurement of magnesium content in tissues

Tissues samples were homogenized and lysed using gentleMACS Dissociators (Catalog No. 130-093-235, Miltenyi Biotec) with RIPA buffer. Subsequently, the tissue lysates were sonicated for 5 min at 227.5 W, with a 3-second on and 10-second off cycle. Bradford protein assay was taken to measure protein concentration, and 350 μL of each sample were taken and used for the measurement of Mg content. Mg content in each tissue was quantified by normalizing to protein concentration. The lyophilized samples were then treated with a suitable amount of 65% nitric acid (HNO_3_) overnight at room temperature. This process was continued by heating the samples in a heating block at 90 °C for about 20 min. Subsequently, an equivalent volume of 30% H_2_O_2_ was added to each sample. The reaction was stopped after an additional 30 min, followed with a further heating step at 70 °C for 15 min. The mean reduced volume was established, and subsequently, the samples were diluted with 1% HNO_3_. Measurements were conducted using an Agilent 7700 series ICP-MS instrument under standard multi-element operating conditions, employing a helium reaction gas cell. Calibration of the instrument was performed using certified multi-element ICP-MS standard calibration solutions with concentrations spanning 0, 5, 10, 50, 100, and 500 ppb for various elements. Moreover, a certified IS solution containing 200 ppb yttrium was employed for an internal control.

### Whole-exome sequencing and data processing

In our previous study, we have utilized a total of 76 paired tumors and corresponding DNTs from CRC patients for WES analysis[35]. The detailed methods have been illustrated in the previous publication. Briefly, genomic DNA was quantified by a Qubit® DNA Assay Kit in Qubit® 2.0 Fluorometer (Catalog No. Q32866, Thermo Fisher Scientific). A total amount of 0.6 µg genomic DNA per sample was used as input material for the DNA sample preparation. Subsequently, WES libraries were prepared and captured using an Agilent SureSelect Human All Exon kit (Catalog No. 5191-5735, Agilent Technologies, Santa Clara, CA). The DNA library, featuring 150 bp paired-end reads, underwent sequencing using an Illumina NovaSeq 6000 System. The initial fluorescence image files acquired from the HiSeq platform were conversed to raw data through base calling and subsequently transformed into FASTQ format. This format includes both sequence information and corresponding sequencing quality details.

### Somatic copy number alteration analysis

The exome sequencing data were initially aligned to the human genome hg19. Following the alignment, copy number variations (CNVs) were detected from the BAM files derived from WES. These CNVs were then utilized for SCNA analysis. To integrate the results obtained from individual patients and identify recurrently amplified or deleted focal genomic regions in our samples, we employed the GISTIC 2.0 software (https://cloud.genepattern.org)[79]. Additionally, the R package “maftools” was used to display Q values less than 0.1. Moreover, to minimize false positives, the related parameters were set as follows: refgene file = Human_Hg19.mat, focal length cutoff = 0.50, gene gistic = yes, confidence level = 0.99. All other parameters were maintained at their default settings.

### Differential abundance analysis

The cleaned proteome and phosphoproteome data underwent a differential abundance analysis between tumors and DNTs through the Wilcoxon rank-sum test. Variables that contain > 50% missing values are excluded. Significance was considered when *P* < 0.05, and fold change (FC) was calculated as the median log_2_(FC). Differential proteins and phosphosites in cell lines were detected through the Student’s *t*-test. Significance was considered when *P* < 0.05, and FC was calculated as the median log_2_(FC). Pathway enrichment analysis was performed based on the DAVID Bioinformatics Resources (https://david.ncifcrf.gov/)[80, 81] and Metascape database (http://metascape.org)[82].

### Correlation analysis

Pearson’s correlation analysis was performed between IS samples or QC samples to assess the quality of the proteome and phosphoproteome data. Mg content and the levels of proteins or phosphosites were calculated by the Spearman’s rank correlation analysis.

### Univariate survival analysis

The optimal cutoff point for the selected samples was calculated by the R package “survminer”. To assess differences between the categorical variables, the log-rank test was applied, and two-tailed tests and *P* values < 0.05 were used for significance evaluation. In the R package “survminer,” survival curves were created through the Kaplan−Meier method for specific variables of interest. The estimation of hazard ratios (HRs) and their 95% confidence intervals (CIs) was performed using the “coxph” function from the R package “Survival.”

### Western blotting analysis

Proteins were extracted from tissues and cells with RIPA buffer and then quantified by the Bradford assay. SDS-PAGE (10%) was used to separate the protein samples, which were then transferred to PVDF membranes (Catalog No. ISEQ00010, Darmstadt, Germany). Following that, PVDF membranes were blocked using 5% milk in PBST and then incubated with an E-cadherin antibody (1:1000; Catalog No. 20874-1-AP, Proteintech, Wuhan, China), N-cadherin antibody (1:1000; Catalog No. 22018-1-AP, Proteintech), Vimentin antibody (1:1000; Catalog No. 10366-1-AP, Proteintech), Vinculin antibody (1:1000; Catalog No. 66305-1-Ig, Proteintech), MMP2 antibody (1:1000; Catalog No. 10373-2-AP, Proteintech), Cingulin antibody (1:1000; Catalog No. 21369-1-AP, Proteintech), DBN1 antibody (1:1000; Catalog No. 10260-1-AP, Proteintech), ACTN4 antibody (1:1000; Catalog No. 19096-1-AP, Proteintech) or GAPDH antibody (1:5000; Catalog No. 60004-1-Ig, Proteintech) overnight at 4 °C. Following overnight incubation, underwent three washes with PBST, followed by incubation with the appropriate secondary antibody at room temperature for 1 hour. Finally, the membrane was subjected to chemiluminescent detection using the Immobilon Western HRP Substrate (Catalog No. WBKLS0500, Sigma).

### Cell culture and generation of stable cell lines

The HCT116 and DLD-1 human CRC cell lines were obtained from the Cell Bank/Stem Cell Bank, Chinese Academy of Sciences. Additionally, HCT116 and DLD-1 cells were cultured in DMEM (Catalog No. C11995-065, Gibco, Grand Island, NY) and RPMI 1640 (Catalog No. 10270-106, Gibco) with 10% FBS (Catalog No. FCS500, ExCell, Shanghai, China), 100 units of penicillin, and 100 μg/mL streptomycin (Catalog No. 15140-122, Gibco), respectively.

To construct DBN1-GFP-tagged and DBN1-Flag-tagged cells, the cDNA of *DBN1* was inserted into the pCDH-LGFP and pCDH-3 ×Flag vectors, respectively. The PLKO.1 vector was used for *DBN1* and *ACTN4* knockdown. To generate cells overexpressing *DBN1* or cells with silenced *DBN1* and *ACTN4*, a co-transfection approach was employed using HEK293T cells. The pSPAX2, pMD2. G, pCDH-LGFP-DBN1, pCDH-3 ×Flag-DBN1, shDBN1, shACTN4, and their respective control plasmids were co-transfected into HEK293T cells. 48 hours later, the medium containing the virus was collected and filtered. To enhance transfection efficiency, 10 mg/mL Polybrene (Catalog No. S2667, Sigma) was added. Subsequently, cells underwent infection and selection with 1 μg/mL puromycin (Catalog No. ST551, Beyotime) for 48 hours. The primer sequences for *DBN1* and *ACTN4* are listed as follows:

shDBN1 # 1: 5’-CCGGCTGTGGAAATGAAGCGGATTACTCGAGTAATCCGCTTCATTTCCACAGTTTTTG-3’

shACTN4 # 1: 5’-CCGGCATCGCTTCCTTCAAGGTCTTCTCGAGAAGACCTTGAAGGAAGCGATGTTTTTG-3’

shACTN4 # 2: 5’-CCGGCCTGTCACCAACCTGAACAATCTCGAGATTGTTCAGGTTGGTGACAGGTTTTTG-3’

The primer sequences for *DBN1*-overexpressing were as follows:

DBN1-PCDH-GFP: 5’-GATTCTAGAGCTAGCGAATTCATGGCCGGCGTCAGCTTCAGC-3’

DBN1-PCDH-Flag: 5’-GATGACAAGTCTAGAGAATTCATGGCCGGCGTCAGCTTCAGC-3’

The primer sequences for *DBN1* mutants were as follows: DBN1^S142A^: 5’-CGCGACTCTCCGCCCCTGTGCTGCA-3’ DBN1^S142D^: 5’-CGCGACTCTCCGACCCTGTGCTGCA-3’

### Cell migration assay

In the cell migration assay, 8.0 mm, 24-well plate chamber inserts were used (Catalog No. 354578, Corning Life Sciences, Corning, NY). In the upper chamber of the inserts, a total of 3 ×10^5^ cells suspended in 200 μL of serum-free medium were added, and 800 μL of 10% FBS medium was added to the bottom chamber. 24 hours later, the cells were fixed with 4% PFA for 15 min and then stained with 0.5% crystal violet for another 15 min. Cells on the upper surface of the inserts were retained, while cells on the underside were gently removed using a cotton swab. The captured images were analyzed using ImageJ software to quantify the number of cells.

### Wound healing assay

HCT116 and DLD-1 cells were plated in a 6-well plate, and a 10 μL pipette tip was used to generate a straight-line wound. The wells were washed with PBS and replenished with serum-free medium. Subsequently, the wounds were photographed at 0 hours and 48 hours after the injury. ImageJ software was utilized to measure the width of the gap.

### Immunofluorescence

Immunofluorescence assays were performed in 96-well plates (Catalog No. 6055300, PerkinElmer, Waltham, MA), and 1 ×10^4^ cells were plated and incubated for 24 hours. Subsequently, the cells were fixed with 4% PFA for 15 min, three washes with PBS, and then permeabilized and blocked with 0.5% Triton X-100 (Catalog No. T8200, Solarbio, Beijing, China) for 5 min. Next, the cells were incubated with Actin-Tracker Red-594 (Catalog No. C2205S, Beyotime) for about 30 min, followed by 4’,6-diamidino-2-phenylindole (Catalog No. C0060, Solarbio) for about 5 min. Images were photographed by Opera Phenix Plus (Catalog No. HH14001000, PerkinElmer), and then quantified by using Harmony software.

### Coimmunoprecipitation

Cells were lysed using RIPA buffer on ice for 30 min, then centrifuged at 20,000 g for 10 min. To pull down DBN1-Flag, anti-Flag magnetic beads (Catalog No. HYK0207, MedChemExpress, Monmouth Junction, NJ) were utilized. The elution of interacting proteins was carried out using 1 × SDS loading buffer (Catalog No. P0015A, Beyotime) and subjected to heating at 95 °C for 5 min. The eluted products were subsequently analyzed either through Western blotting or LC-MS/MS.

## Ethical statement

Written informed consents and approvals for all tissue specimens were obtained from the Research Ethics Committee (Permission number: 2020 (374)).

## Data availability

The MS proteomics data of CRC have been deposited to the ProteomeXchange Consortium via the iProX partner repository with the dataset identifier PXD039360. The mass spectrometry phosphoproteomics data of CRC and cells have been deposited in the ProteomeXchange Consortium and are available using the iProX accession number PXD042746. The raw GSE sequence data reported in this paper have been deposited in the Genome Sequence Archive[83] in the National Genomics Data Center, China National Center for Bioinformation/Beijing Institute of Genomics, Chinese Academy of Sciences (GSA-Human: HRA003386), which are publicly accessible at https://ngdc.cncb.ac.cn/gsa-human.

## Code availability

The source code is freely available at the National Genomics Data Center (NGDC) BioCode (https://ngdc.cncb.ac.cn/biocode/tools/BT007400).

## CRediT authorship contribution statement

**Rou Zhang:** Investigation, Validation, Visualization, Writing– original draft. **Meng Hu:** Software, Formal analysis, Visualization, Writing – original draft. **Yu Liu:** Investigation, Validation, Visualization. **Wanmeng Li:** Visualization. **Zhiqiang Xu:** Investigation. **Siyu He:** Visualization. **Ying Lu:** Visualization. **Yanqiu Gong:** Validation. **Xiuxuan Wang:** Validation. **Shan Hai**: Resources, Funding acquisition. **Shuangqing Li**: Resources, Funding acquisition. **Shiqian Qi**: Supervision. **Yuan Li:** Resources. **Yang Shu:** Data curation, Software. **Dan Du**: Resources. **Huiyuan Zhang**: Resources. **Heng Xu:** Resources, Funding acquisition. **Zongguang Zhou:** Resources, Funding acquisition. **Peng Lei:** Resources, Project administration, Conceptualization. **Hai-Ning Chen:** Resources, Writing – review & editing, Supervision. **Lunzhi Dai:** Conceptualization, Writing – review & editing, Supervision, Project administration, Funding acquisition. All authors have read and approved the final manuscript.

## Acknowledgments

This work was supported by the National Key R&D Program of China (No. 2022YFA1303200), National Natural Science Foundation of China (No. 82073221, 31870826, and 82073246), Sichuan Science and Technology Project (2021YFS0134), National Clinical Research Center for Geriatrics of West China Hospital (No. Z2021JC005) and the 135 project of West China Hospital (No. ZYYC23013, ZYYC23025, ZYGD20006, and 2016105).

## ORCID

ORCID 0009-0001-5056-8345 (Rou Zhang)

ORCID 0000-0001-5113-2888 (Meng Hu)

ORCID 0000-0001-5359-1906 (Yu Liu)

ORCID 0009-0009-1357-3629 (Wanmeng Li)

ORCID 0000-0002-8133-1228 (Zhiqiang Xu)

ORCID 0009-0007-4350-4232 (Siyu He)

ORCID 0000-0002-6424-5424 (Ying Lu)

ORCID 0000-0001-7890-5158 (Yanqiu Gong)

ORCID 0000-0002-4925-7048 (Xiuxuan Wang)

ORCID 0009-0007-7718-4249 (Shan Hai)

ORCID 0000-0001-8510-7568 (Shuangqing Li)

ORCID 0000-0002-7589-8877 (Shiqian Qi)

ORCID 0000-0003-0274-855X (Yuan Li)

ORCID 0000-0002-8304-1531 (Yang Shu)

ORCID 0000-0001-9745-8197 (Dan Du)

ORCID 0000-0002-1532-2462 (Huiyuan Zhang)

ORCID 0000-0002-7748-2621 (Heng Xu)

ORCID 0000-0002-7616-1199 (Zongguang Zhou)

ORCID 0000-0001-5652-1962 (Peng Lei)

ORCID 0000-0003-0104-8498 (Hai-Ning Chen)

ORCID 0000-0002-3003-8910 (Lunzhi Dai)

## Supplementary material

### Supplementary Figure Legends

**Figure S1.**
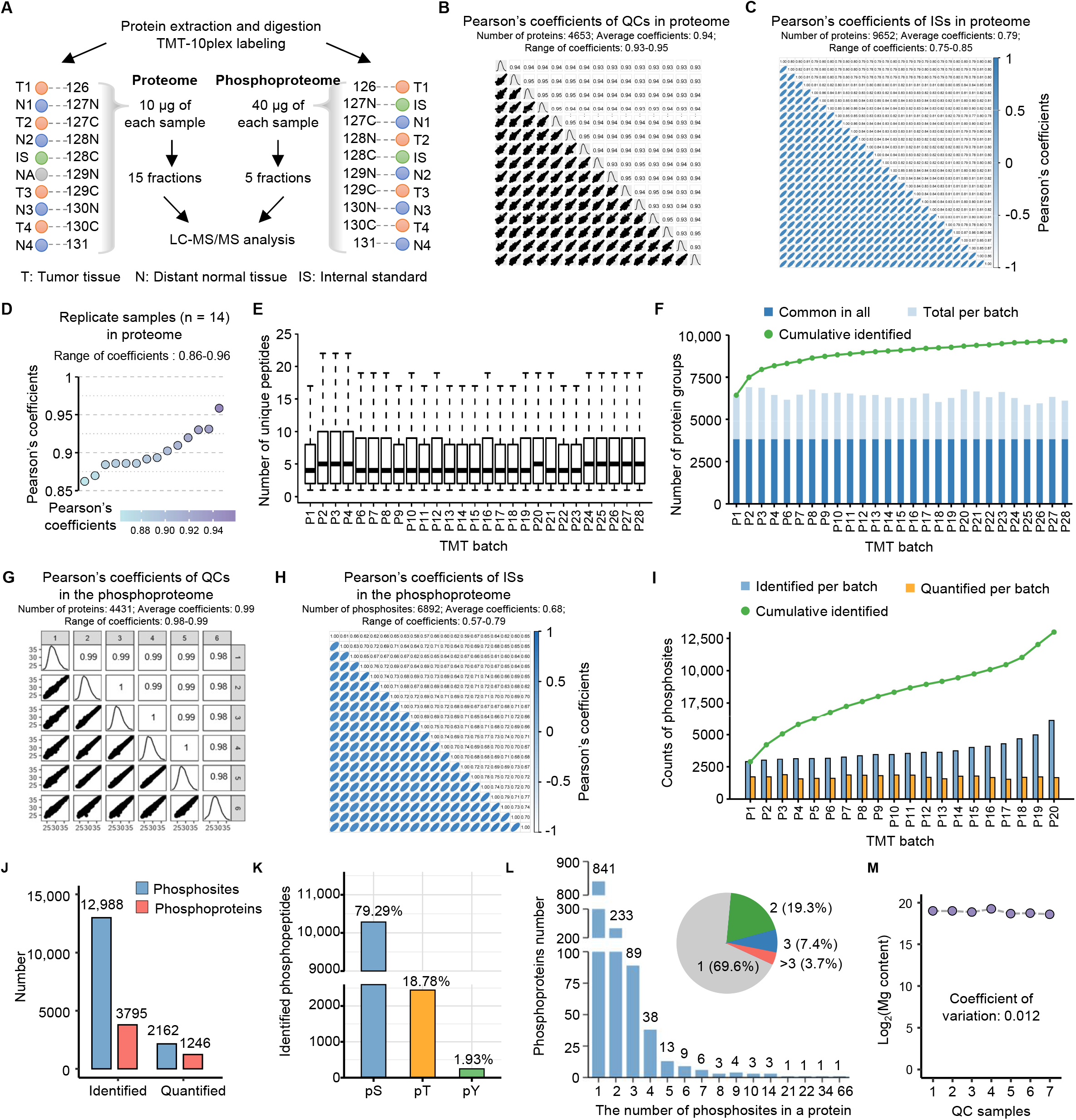
Quality control analysis of proteome and phosphoproteome data and Mg content detection. **A.** Workflow of proteome and phosphoproteome experiments. **B.** Pearson’s correlation analysis of 15 QC samples of the proteome to evaluate machine stability. A scatterplot matrix was generated for pairwise calculation of Pearson’s correlation coefficients among the samples, with density plots on the diagonal and correlation values displayed in the upper triangle. **C.** Pearson’s correlation analysis of 27 internal standards to evaluate the robustness of TMT-based quantification in the proteomic study. Top right: pairwise calculation of Pearson’s correlation coefficients among the 27 ISs. Bottom left: elliptic chart shows pairwise comparison of the 27 ISs. **D.** Pearson’s correlation analysis of replicate samples. **E.** Distribution of unique peptides among 27 TMT-labeled batches. **F.** Protein counts of each batch (light blue), counts of common proteins (blue) and cumulative protein counts among the 27 batches. **G.** Pearson’s correlation analysis of 6 QC samples of the phosphoproteome to evaluate machine stability. A scatterplot matrix was generated for pairwise calculation of Pearson’s correlation coefficients among the samples, with density plots on the diagonal and correlation values displayed in the upper triangle. **H.** Pearson’s correlation analysis of 20 ISs to evaluate the robustness of TMT-based quantification in the phosphoproteomic study. Top-right half: pairwise calculation of Pearson’s correlation coefficients among the 20 ISs. Bottom-left half: elliptic chart shows pairwise comparison of the 20 ISs. **I.** Identified (blue) and quantified (orange) phosphoprotein counts of each batch and cumulative phosphoprotein counts among 20 batches. **J.** Number of identified and quantified phosphosites (blue) and phosphoproteins (red). **K.** Localization and distribution of phosphosites. **L.** Statistics of the number of phosphosites in a protein. **M.** Coefficient of variation of Mg content for the QC samples.

**Figure S2.**
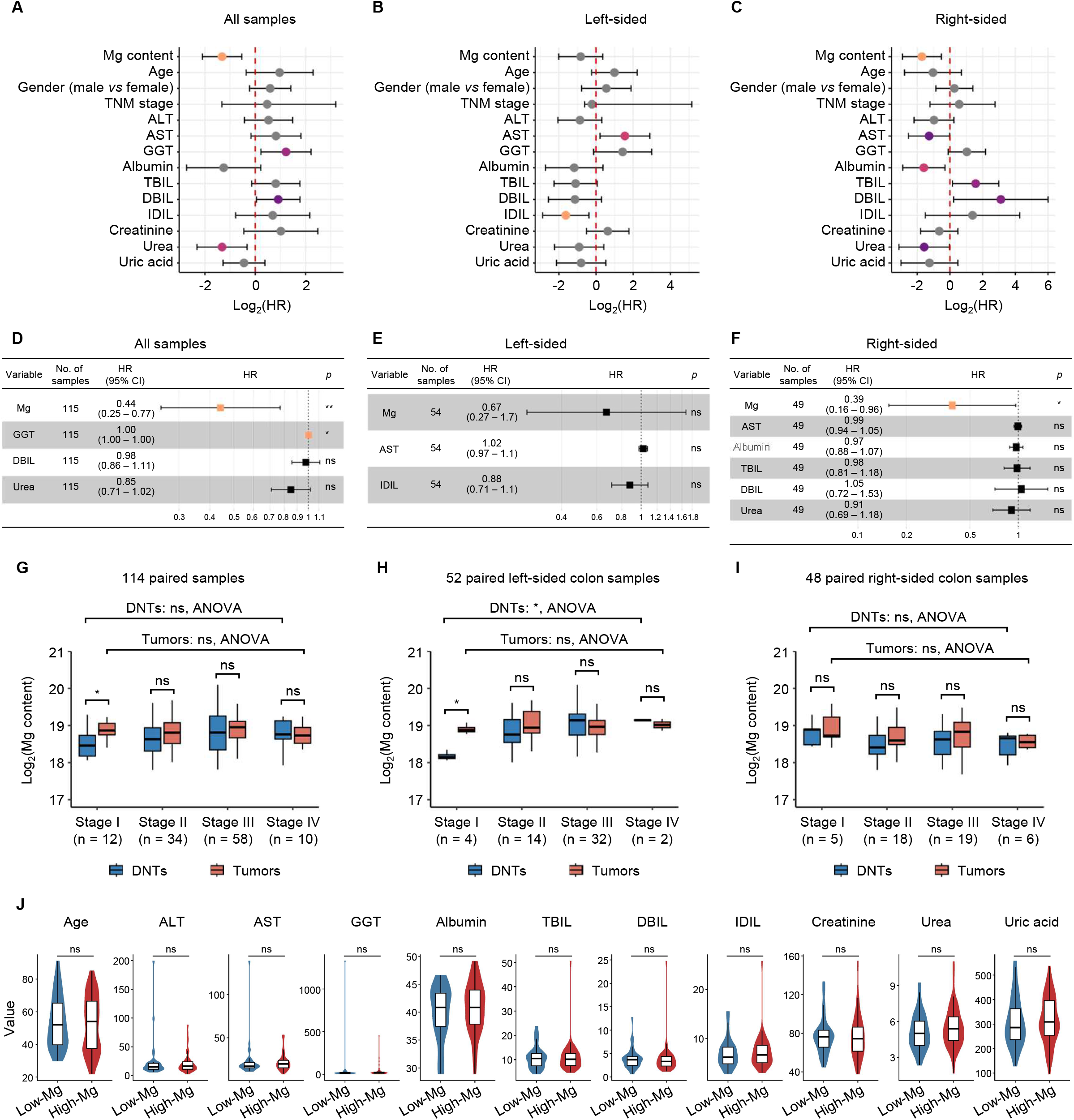
Association analysis between Mg content and TNM stage (or overall survival) **A.–C.** Forest plots of univariable cox models for clinical information with CRC patients in all samples (A), or left-sided (B) or right-sided (C) colon samples. Data are presented as hazard ratio (HR) ±95% confidence interval. **D.**−**F.** Cox multivariable regression models designed to test for prognostic factors. Low Mg content was an independent predictor of outcome in all CRC patients (D) and right-sided CRC patients (E) but not left-sided CRC patients (F). Data are presented as hazard ratio (HR) ±95% confidence interval. **G.**−**I.** Mg content among four TNM stages in tumors and DNTs of all samples (G) or left-sided (H) or right-sided (I) colon samples. **J.** Violin plots illustrating the levels of clinical information, encompassing variables such as age, ALT, AST, etc., for both the High-Mg and Low-Mg groups.

**Figure S3.**
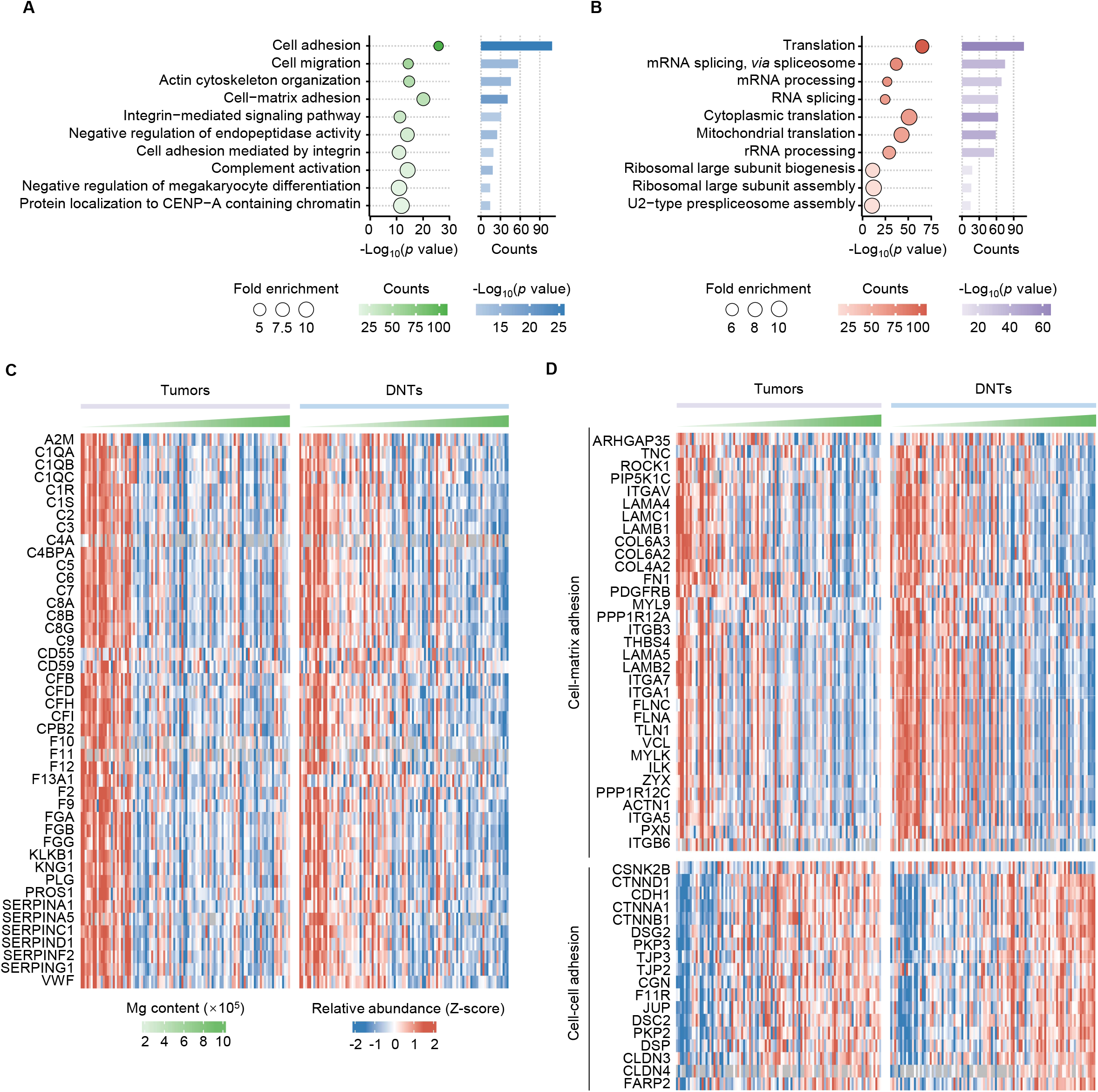
Analysis of Mg-associated proteome and phosphoproteome. **A.** GOBP term enrichment analysis of proteins negatively associated with Mg content using DAVID database. **B.** GOBP term enrichment analysis of proteins positively associated with Mg content using DAVID database. **C.** The expression of complement-associated proteins in the tumors and paired DNTs. **D.** The expression of cell adhesion-associated proteins in the tumors and paired DNTs. GOBP, Gene ontology biological process.

**Figure S4.**
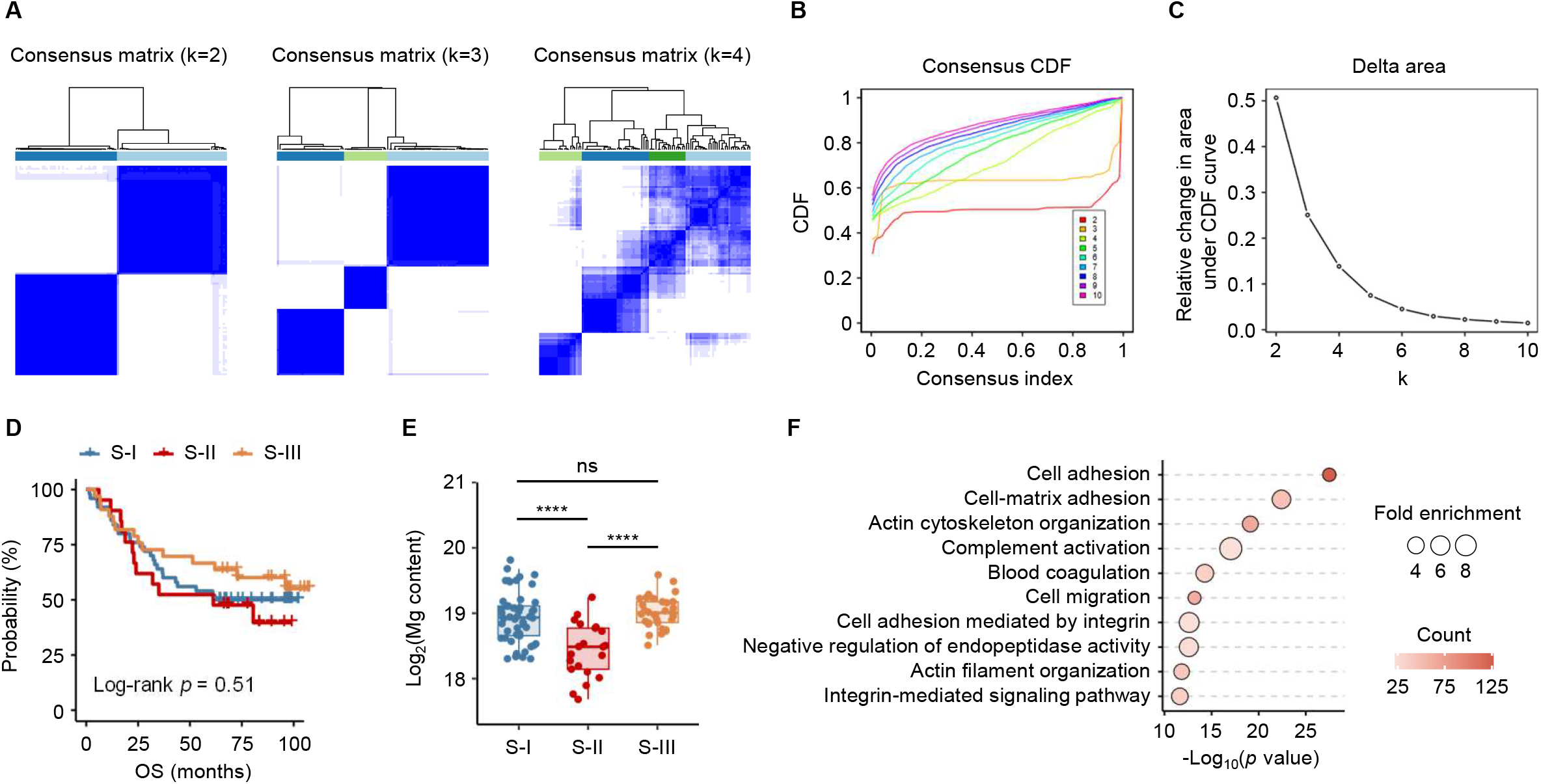
Proteomic classification of CRC tumors. **A.** Consensus matrix of unsupervised clustering based on the top 25 most variable proteins (k = 2, k =3 and k = 4). **B.** The consensus CDF of unsupervised clustering based on the top 25 most variable proteins. **C.** Delta area (change in CDF area) plots for 2-10 clusters. **D.** Kaplan–Meier curves of OS for each proteomic subtype. **E.** Boxplot showing the difference of Mg content among proteomic subtypes. **F.** GOBP enrichment analysis of proteins upregulated in subtype II (subtype II *vs* subtype I and subtype III) using DAVID.

**Figure S5.**
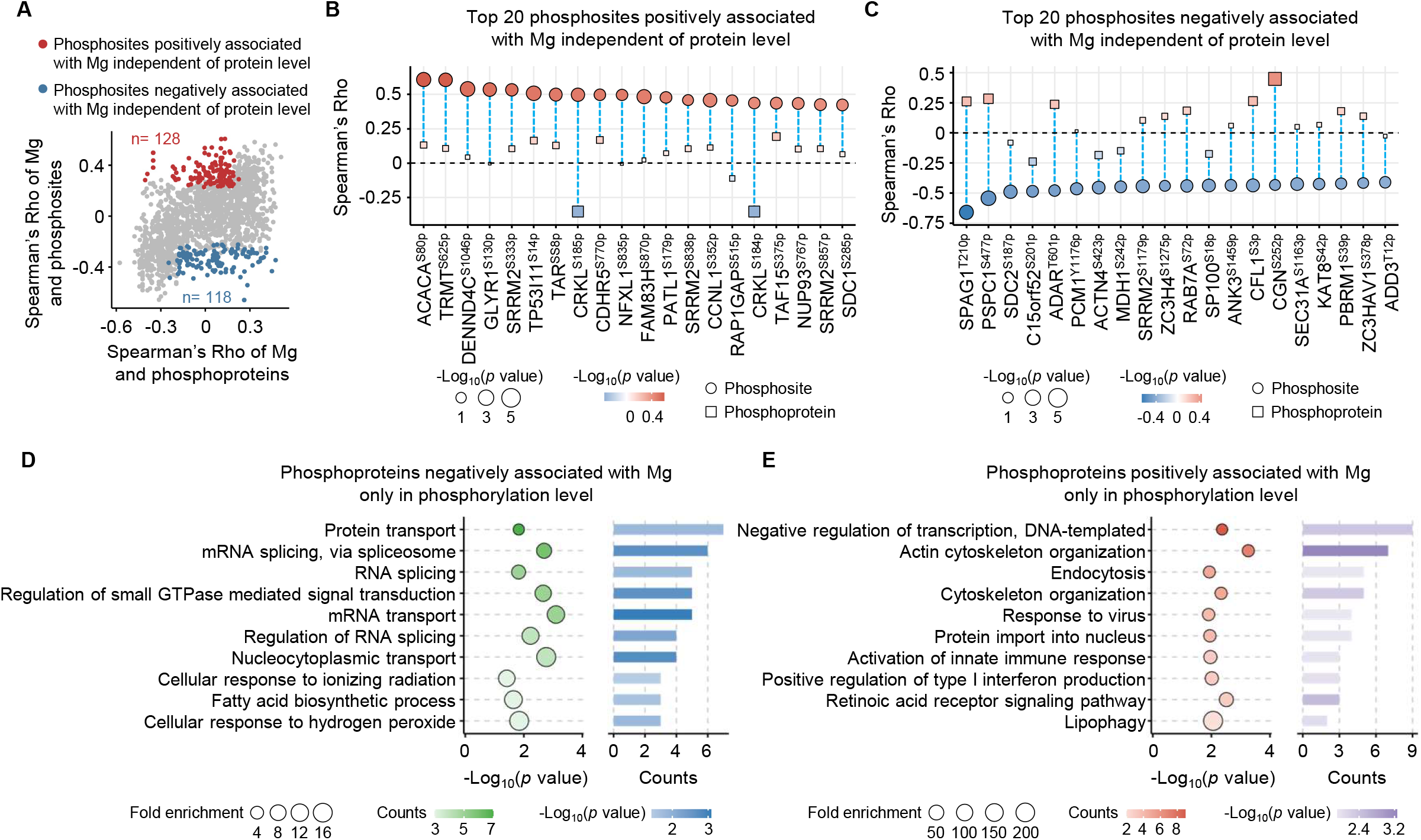
Integrative analysis of proteome and phosphoproteome data. **A.** Comparison between Mg-protein correlations and their Mg-phosphosite correlations. *P* < 0.05. Spearman’s rank correlation test. **B.** and **C.** Top 20 phosphosites positively (B) or negatively (C) associated with Mg content independent of protein level. **D.** GOBP enrichment analysis of phosphoproteins with negatively Mg-associated phosphosites. **E.** GOBP enrichment analysis of phosphoproteins with positively Mg-associated phosphosites.

**Figure S6.**
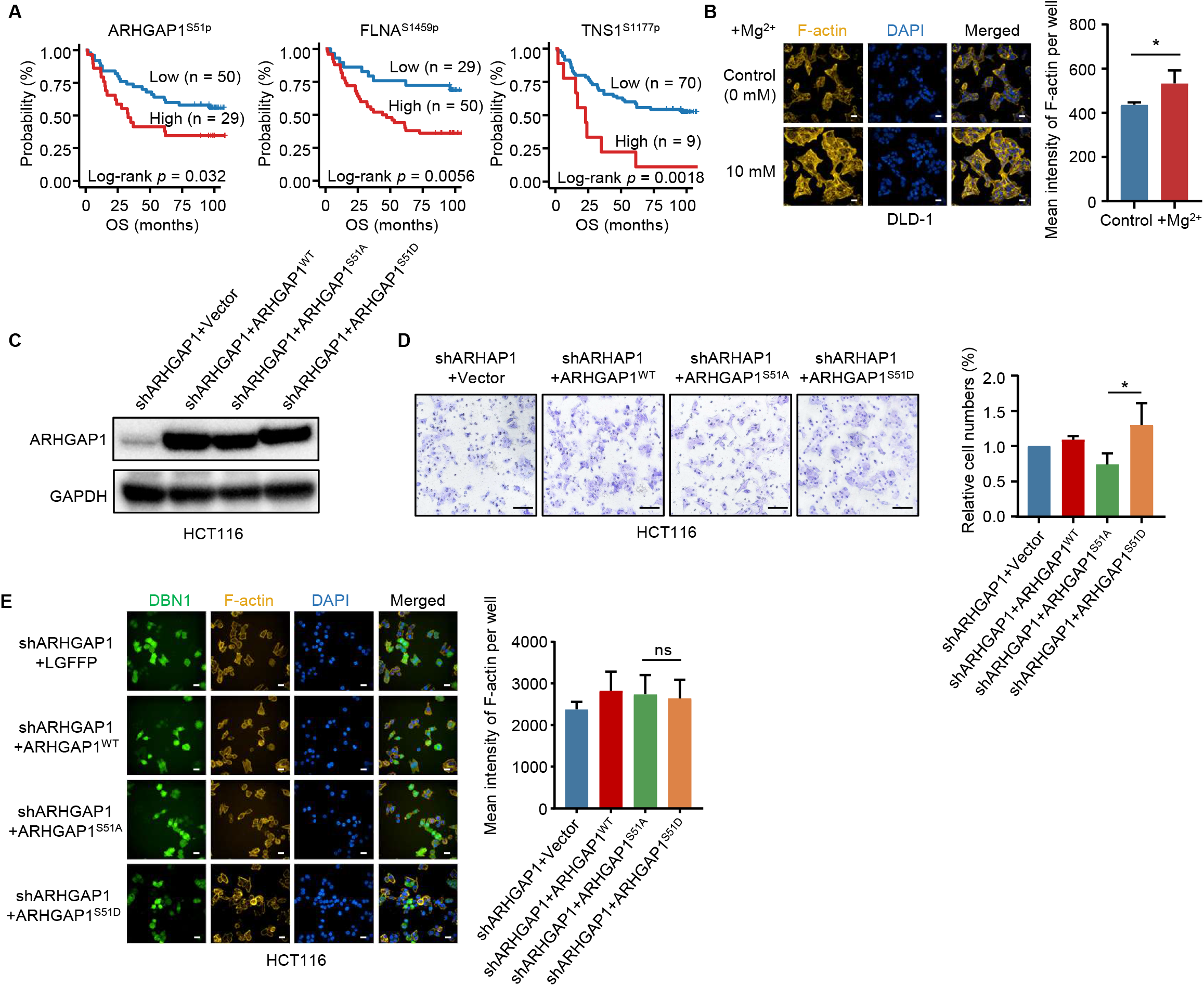
Mg-regulated phosphosites related to cell adhesion. **A.** Survival analysis of 79 CRC patients with different ARHGAP1^S51p^, FLNA^S1459p^ and TNS1^S1177p^ levels in tumors. Log-rank test. **B.** Effects of MgCl_2_ treatment on the formation of F-actin determined by immunofluorescence assays in DLD-1 cells. Scale bars, 500 μm. *, *P* < 0.05, Student’s *t* test. **C.** Immunoblotting assays determined the expression of ARHGAP1 mutations. *, *P* < 0.05, Student’s *t* test. **D.** Transwell assays were performed in cells transfected with empty vector, ARHGAP1^WT^, ARHGAP1^S51A^ and ARHGAP1^S51D^, respectively. Scale bars, 500 μm. **, *P* < 0.01; *, *P* < 0.05. Student’s *t* test. **E.** Effects of empty vector, ARHGAP1^WT^, ARHGAP1^S51A^ and ARHGAP1^S51D^ on F-actin formation determined by immunofluorescence assays. Scale bars, 500 μm. *, *P* < 0.05. Student’s *t* test.

**Figure S7.**
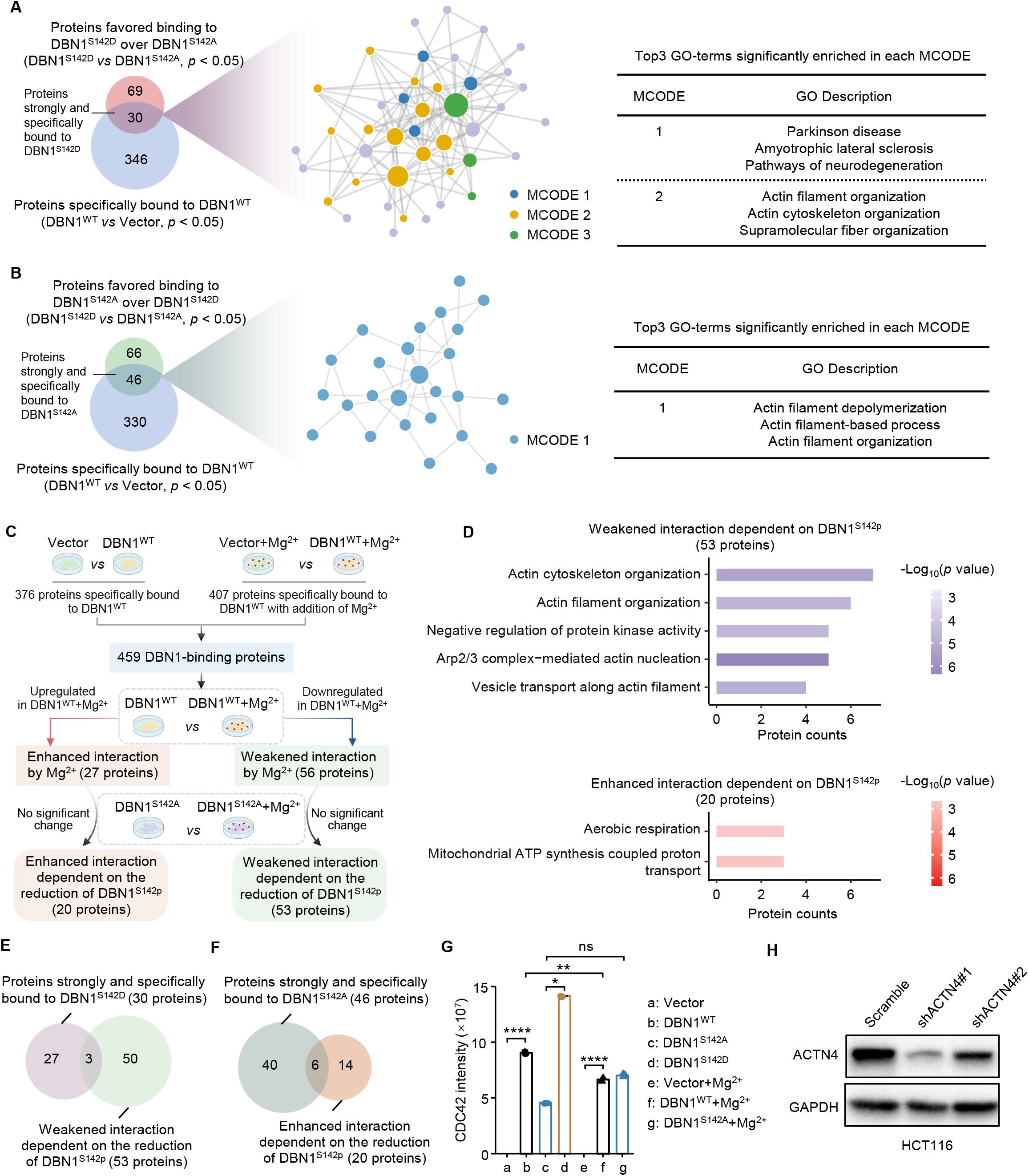
The DBN1-binding proteins affected by DBN1^S142p^ and Mg^2+^. **A. and B.** Venn diagram showing the number of proteins strongly and specifically bound to DBN1^S142D^ (A) and DBN1^S142A^ (B), respectively. Fold changes of proteins between groups > 1.2, *P* < 0.05. Student’s *t* test. Gene annotation enrichment analysis of the proteins obtained in protein-protein interaction network analysis. **C.** Schematic of screening Mg-regulated DBN1 binding proteins affected by DBN1^S142p^. **D.** GO biological process enrichment analysis of Mg-regulated DBN1 binding proteins. **E.** and **F.** Venn diagram showing the number of interacting proteins affected by both DBN1^S142p^ and Mg^2+^. **G.** The levels of the DBN1-binding protein CDC42 under different treatment and immunoprecipitation conditions. ****, *P* < 0.0001; **, *P* < 0.01; *, *P* < 0.05; ns means no significance. Student’s *t* test. **H.** Immunoblots showing the expression of ACTN4.

**Figure S8.**
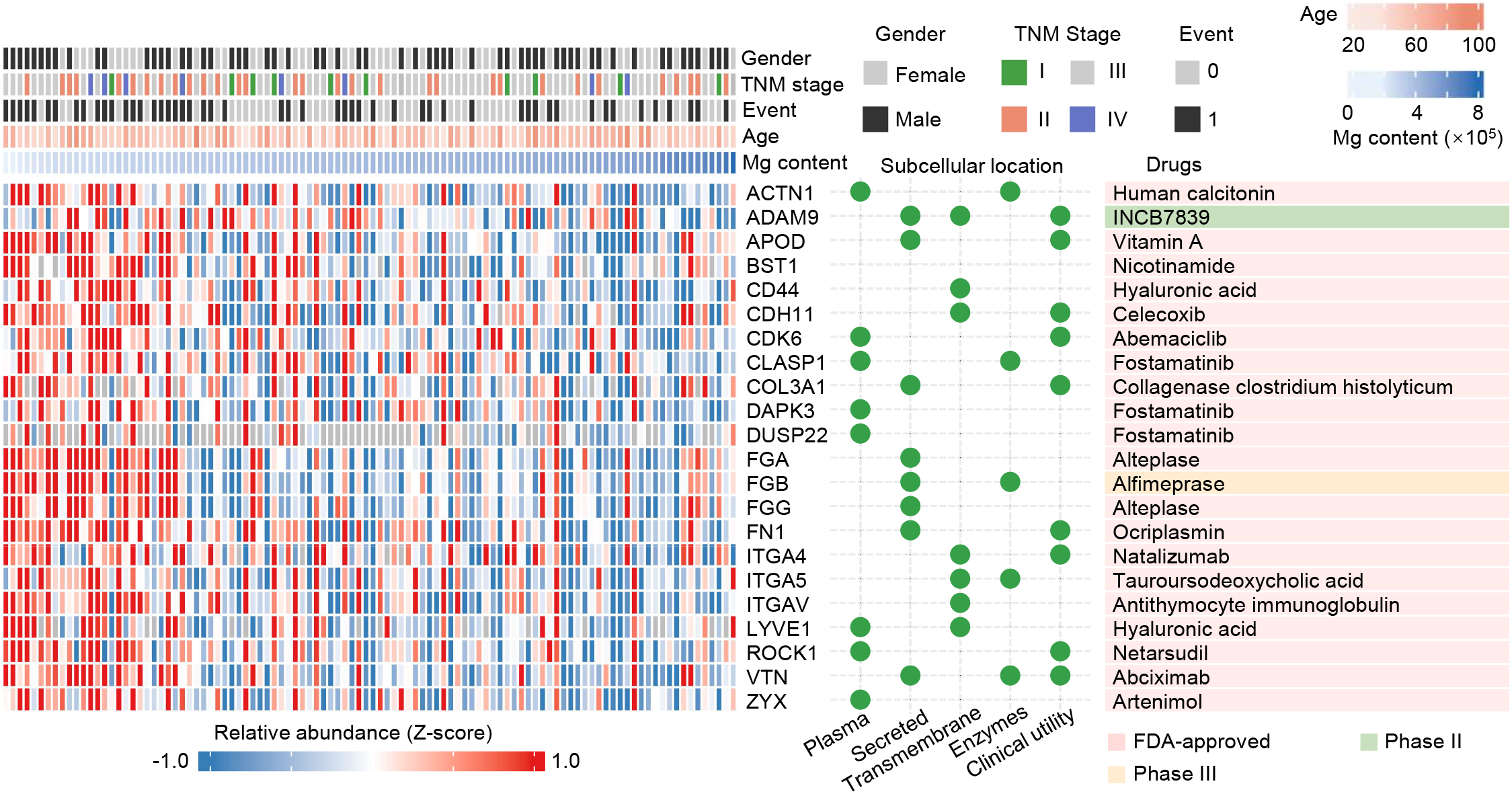
Mg-associated cell-matrix adhesion proteins as drug targets. Cell-matrix adhesion-associated proteins with clinical drugs were negatively associated with Mg content. Spearman’s rank correlation analysis, *P* < 0.05 indicating a significant correlation. The gender, TNM stage, survival event, age and Mg content are annotated above the heatmap. The heatmap depicts the intensity of proteins using the z-score. The corresponding subcellular location (middle) and clinical drugs (right) are listed.

### Supplementary Table Legends

**Table S1. Clinical information of CRC patients and Mg content in tumors and DNTs**.

**Table S2. Genomic data of CRC**.

**Table S3. Proteomics data of CRC**.

**Table S4. Spearman’s rank correlation analysis of the levels of the cell adhesion-related proteins and Mg content.**

**Table S5. Phosphoproteomics data of CRC**.

**Table S6. Phosphoproteome in Mg-treated HCT116 cells**.

**Table S7. Proteins strongly and specifically bound to DBN1 and affected by DBN1^S142p^ and Mg^2+^**.

**Table S8. The subcellular location and drugs of targets.**

